# Context-dependence of T-loop mediated long-range RNA tertiary interactions

**DOI:** 10.1101/2022.12.26.521914

**Authors:** Lisa N. Hansen, Otto A. Kletzien, Marcus Urquijo, Logan T. Schwanz, Robert T. Batey

**Affiliations:** Department of Biochemistry, University of Colorado, Boulder, CO, 80309-0596, USA

**Keywords:** RNA structure, structural motif, regulatory RNA, tertiary interaction, T-loop

## Abstract

The architecture and folding of complex RNAs is governed by a limited set of highly recurrent structural motifs that form long-range tertiary interactions. One of these motifs is the T-loop, which was first identified in tRNA but is broadly distributed across biological RNAs. While the T-loop has been examined in detail in different biological contexts, the various receptors that it interacts with are not as well defined. In this study, we use a cell-based genetic screen in concert with bioinformatic analysis to examine three different, but related, T-loop receptor motifs found in the flavin mononucleotide (FMN) and cobalamin (Cbl) riboswitches. As a host for different T-loop receptors, we employed the *env*8 class-II Cbl riboswitch, an RNA that uses two T-loop motifs for both folding and supporting the ligand binding pocket. A set of libraries was created in which select nucleotides that participate in the T-loop/T-loop receptor (TL/TLR) interaction were fully randomized. Library members were screened for their ability to support Cbl-dependent expression of a reporter gene. While T-loops appear to be variable in sequence, we find that the functional sequence space is more restricted in the Cbl riboswitch, suggesting that TL/TLR interactions are context dependent. Our data reveal clear sequence signatures for the different types of receptor motifs that align with phylogenic analysis of these motifs in the FMN and Cbl riboswitches. Finally, our data suggest the functional contribution of various nucleobase-mediated long-range interactions within the riboswitch subclass of TL/TLR interactions that are distinct from those found in other RNAs.

**Highlights:**

- The T-loop motif frequently mediates tertiary interactions in RNA
- Activity-based screen used to explore T-loop mediate interactions in riboswitches
- Results from the screen were consistent with phylogenetic analysis
- T-loop/T-loop recepto­­­­r interactions are context dependent

## Introduction

Advances in understanding the three-dimensional (3D) structure of RNA over the last several decades reveal that one of the central organizing principles of higher order architecture is the use of highly recurrent structural motifs to promote helical packing and tertiary structure.^1, 2^ From analysis of large RNAs, a set of these motifs have been identified, such as tetraloops and their receptors, kink-turns, loop E motifs, kissing loops, and pseudoknots.^3–6^ A detailed understanding of the folding and structural properties of these recurrent motifs is crucial for advancing our knowledge of RNA function and design. In the current view of RNA folding, secondary structure (i.e., base paired helices) forms rapidly and is followed by the slower process of tertiary structure acquisition, which is mediated by structural motifs that promote long-range interactions.^7, 8^ These modules are key control centers for establishing the folding pathway and rate of folding of larger RNAs, and small changes in the sequence of these motifs can dramatically alter their folding behavior.^9^ Recurrent RNA modules serve as building blocks used in the modeling of RNA structure and design of novel functional RNAs.^2, 10–12^ One premise involved in the design of functional 3D RNA structures is the modularity and portability of these elements such that they are discrete modules that can be incorporated into different structural contexts while maintaining function.^2, 13, 14^

T-loops are one of the recurrent structural motifs that RNAs use to establish higher order architecture and mediate intermolecular interactions.^15^ This module, first identified in the third loop of tRNA that is referred to as the tRNA T-loop,^16, 17^ is observed in mRNAs including riboswitches^18–24^ and T-box elements,^25, 26^ the ribosome,^27–30^ transfer-messenger RNA,^31^ ribonuclease P RNA,^32, 33^ group II self-splicing introns,^34^ a small ribozyme,^35^ viral RNAs that mimic tRNA structure,^36^ bacterial Y RNA^37^, and a synthetic aptamer.^38^ Within these RNAs, the T-loop is observed to promote long-range tertiary interactions as in the T-loop/D-loop interaction of tRNA,^16, 17^ intermolecular RNA-RNA interactions such as observed in the T-box/tRNA interaction,^25, 26^ and small molecule recognition in the TPP riboswitch and the 5-hydroxytryptophan aptamer.^22–24, 38^ Finally, in the ribosome, ribosomal proteins are observed interacting with many of the T-loop motifs.^28, 29^ The dispersal of the T-loop in biology likely arises from its ability to broadly mediate intra- and intermolecular interactions across disparate structural contexts.

The T-loop is defined as a five-nucleotide structural motif (**Figure 1(A)**).^15^ The closing base pair of the loop is often a non-canonical (e.g., reverse Hoogsteen) pyrimidine-adenosine (Y-A) base pair, but the module can accommodate a variety of base pairings between positions 1-5. Following the numbering in Figure 1(A), nucleobase 2 stacks on the that of 1 and is positioned to form a hydrogen bond with the ribose/phosphate of nucleotide 5. Nucleobase 4 can either interact with nucleobase 2 (**Figure 1(A)**) or provide a stacking interaction with a nucleotide from elsewhere in the RNA or a ligand. Finally, nucleotide 3 is available to form a long-range base-base interaction, such as the essential G19-C56 base pair between the T-loop and D-loop of tRNA.^43, 44^ A recent crystal structure of a hairpin with an isolated T-loop demonstrates that its canonical structure can be adopted in the absence of interacting partners, indicating that the fold is intrinsic to a set of sequences in the context of a terminal or internal loop motif.^45^

**Figure 1.**
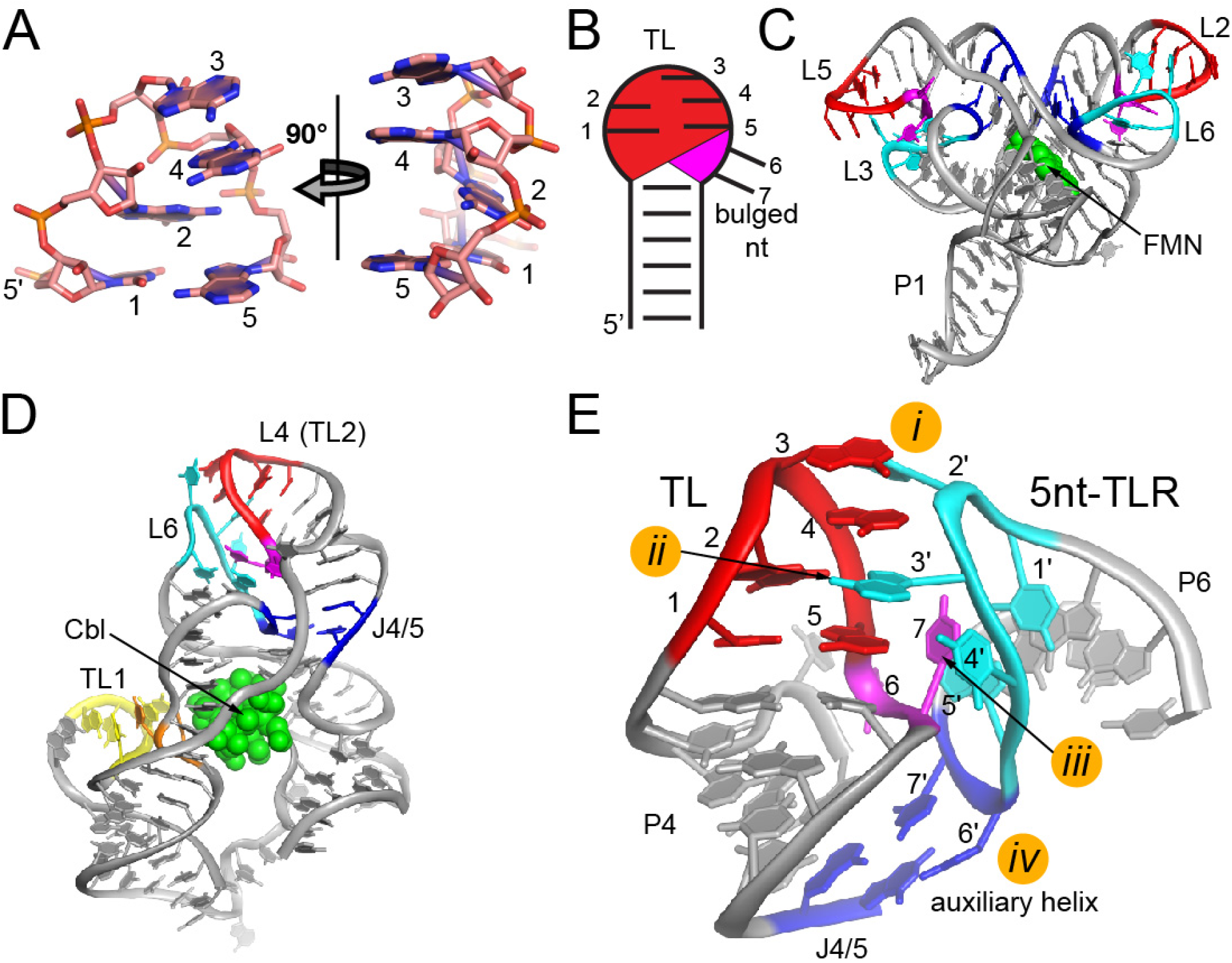
Structure of the FMN/Cbl TL/TLR interaction. (A) Structure of the classic T-loop (TL) motif showing the nucleobase configuration of the five nucleotides (positions 1-5). Panel on the left represents a front view emphasizing all nucleobases and the right is a 90° clockwise rotation of the left view emphasizing the stacking interactions and the space between positions 4 and 5. (B) Schematic of the terminal loop in FMN/Cbl riboswitches harboring the T-loop motif (red) and two bulged nucleotides (magenta). (C) Cartoon representation of the global architecture of the FMN riboswitch bound to FMN (green) (PDB 3F2Q). The T-loop motifs are shown in red, the two nucleotides 3′ of the TL in the terminal loop are magenta, the T-loop receptor is colored cyan, and th^′^e two nucleotides 3′ immediately 3′ of the TLR in the terminal loop are dark blue along with two nucleotides of the helix of the T-loop stem-loop that form a two-nucleotide helix (auxiliary helix). This color scheme will be used throughout this work. (D) Cartoon representation of the *env*8 Cbl riboswitch bound to Cbl (green). The TL/5nt-TLR interaction is shown in the same colors as in panel C, with a second T-loop motif and two interacting nucleotides that support the ligand binding pocket are shown in yellow and orange, respectively (PDB 4FRN). (E) Closeup view of the TL/5nt-TLR interaction in the *env*8 aptamer (PDB 4FRG). Color scheme is the same as in panels C and D. The four nucleobase-mediated contacts facilitated by the T-loop and the 5nt-TLR are labeled as *i*-*iv*.

While the T-loop has been examined in detail in different biological contexts,^15, 40, 41^ the various receptors that it interacts with are not as well defined. The most well established is the D-loop of tRNA, interacting with the T-loop to form the elbow of the tRNA.^46^ Several widely distributed classes of riboswitches^47, 48^ contain T-loop-mediated interactions central to their 3D architecture, including the flavin mononucleotide (FMN)^21^ and class-I^19, 20^ and class-II cobalamin (Cbl) riboswitches.^18, 19^ These RNAs have universally conserved T-loop motifs embedded in terminal loops in which the T-loop comprises the first five nucleotides on the 5′ -side of the loop followed by two bulged nucleotides on the 3′ -side of the loop (**Figure 1(B)**). The FMN riboswitch has a symmetric architecture in which T-loops interact with a loop motif that will be referred to as the T-loop receptor (TLR).^21, 49^ In the crystal structure of the *Fusobacterium nucleatum* FMN riboswitch the terminal loops of helices P2 and P6 (L2 and L6, respectively) form one interaction and those of P3 and P5 (L3 and L5, respectively) form the second (red and cyan; **Figure 1(C)**).^21^ Across all Cbl riboswitches, terminal loop L4 comprises a sequence and structure consistent with a T-loop motif.^19, 20, 50, 51^ This loop interacts with either a terminal loop (class-II, **Figure 1(D)**) or internal loop (class-I and class-II) element to facilitate organization of an adjacent four-way junction formed by P3-P6 that is the core of the Cbl binding site.^19, 20^ Given that L4 is extensively protected from chemical probing agents in the unbound state,^19, 50, 52^ organization of the TL/TLR interaction does not require ligand binding.

Crystal structures of FMN and Cbl riboswitches reveal that the TL/TLR interaction common to these RNAs involves a set of specific interactions between sub-elements of the loops. While the T-loop motif is a five-nucleotide sequence, the FMN/Cbl TL/TLR motif’s L4 is always seven nucleotides.^47, 48^ Nucleotides 1-5 form the T-loop and interact with two nucleotides donated by the TLR (contacts *i* and *ii*, **Figure 1(E)**). Contact *ii*, where a nucleotide from the TLR intercalates between position 4 and 5 of the T-loop (**Figure 1(E)**), is nearly universal amongst the T-loop-mediated interactions. The third contact involves nucleotides 6 and 7 of the terminal loop, where the T-loop resides. These nucleotides are flipped out and nucleotide 7 forms hydrogen bonds to the TLR (contact *iii*, **Figure 1(E)**). Finally, the TLR has two bulged nucleotides that base pair with nucleotides immediately 3′ of the stem of P4, which we refer to as the “auxiliary helix” in this work (contact *iv*, **Figure 1(E)**). Together, these four interactions mediated by sub-elements of the TL and TLR serve to establish the full long-range TL/TLR interaction, although the relative importance of these contacts to the stability of the interaction is not known.

The FMN/Cbl TLR motif is observed in three distinct loops. The first two are six and seven nucleotide terminal loops, which we call the 4nt-TLR and 5nt-TLR, respectively (the rationale for this nomenclature is that, like the T-loop, the last two nucleotides of the TLR loops are bulged out). Comparison of the structure of the two loops (L6 of the *F. nucleatum* FMN riboswitch^21^ and L6 of *env*8 Cbl riboswitch^19^) indicates that the 4nt-TLR has a deletion of position 2 of the 5nt-TLR. The six and seven nucleotide terminal loop motifs are found broadly in the FMN and class-II Cbl riboswitches. The third motif is an internal loop motif (referred to as the IL-TLR) that is found across FMN and class-I and class-II Cbl riboswitches.^47, 48^ The three motifs appear to be interchangeable, as all three types of FMN/Cbl TLR are found mediating this interaction at equivalent sites in the FMN and class-II Cbl riboswitches.

Riboswitches, a small-molecule dependent mRNA regulatory element,^53, 54^ are an ideal model system for developing a functional understanding of RNA structures. For example, the SAM-I riboswitch has been extensively used to explore structural and functional diversity of the kink-turn motif.^55–58^ In those studies, the riboswitch served as a host for various kink-turn motifs that were analyzed using a combination of X-ray crystallography and biophysical approaches. More recently, a genetic screen was employed to study a T-loop motif observed to buttress the ligand binding pocket of the class-II *env*8 Cbl riboswitch.^59^ This approach uses the ability of the riboswitch to regulate expression of a reporter gene to assess the activity of different variants. The *env*8 Cbl riboswitch is an OFF switch in which the binding of a ligand to the mRNA stabilizes formation of a structure that blocks access of the ribosome to the message.^19^ Thus, the ability of a sequence variant observed in the genetic screen to repress gene expression is a function of the ability of the RNA to rapidly acquire an architecture that organizes the Cbl binding pocket, bind the effector ligand, and form the structure that represses translation within a short kinetic window.^60^ Since the TL/TLR interaction of Cbl riboswitches is the key tertiary interaction involved in organization of the binding pocket, variation in this interaction will likely have a strong influence on the kinetics of folding and therefore its regulatory function. As such, this model system allows us to explore the sequence space that can support a particular RNA structure or tertiary interaction and assess the performance of the variants emerging from the screen to determine how sequence variation with the TL/TLR interaction influences function.

In this study, we have applied a functional assay, previously used to assess structure-function relationships in the class-II *env*8 Cbl riboswitch,^19, 59–62^ to assess the sequence diversity supporting the three types of FMN/Cbl TL/TLR interactions and the impact of that diversity on regulatory function. To accomplish this, we separated different components of the TL/TLR interactions into libraries in which the nucleotides within each site were fully randomized and subjected to a screen to find functional variants within that library. Variants identified from the screen were assessed for their ability to regulate expression of a fluorescent protein reporter in a cellular context. This analysis enabled us to probe regions of T-loop and TLR sequence space that were highly functional and compare them to the sequence space of these motifs observed in biological FMN and Cbl riboswitches. Analysis of a library encompassing the T-loop and two directly interacting nucleotides of the 5nt-TLR reinforce a prior structure-based analysis^15^ that sequences supporting the T-loop of the FMN/Cbl riboswitches are distinct from those found in tRNAs but similar to other large RNAs. Findings resulting from screens of the three distinct TLRs reveal clear patterns of sequence preferences that impact the regulatory function of the riboswitch. These preferences establish the importance of key positions in each type of TLR for activity and point to key differences between them that may establish their differential use in biology.

## Results and Discussion

### TLRs are context-dependent in FMN and Cbl riboswitches

As a basis of comparison of functional sequences that emerge from the genetic screen, we first sought to create a structure-based alignment of TL/TLR interactions from the FMN and Cbl riboswitch families. To this end, the family sequence alignments of the FMN and class-I and class-II Cbl riboswitches were taken from Rfam 14.0^63, 64^, manually curated using JalView^65, 66^, and aligned in MUSCLE^67, 68^. The aim of this was to create a set of high-confidence alignments of each family in which individuals not containing all the conserved sequences necessary to support both folding and ligand binding were removed, as guided by representative crystal structures. Sequences within the new alignments were further separated into three groups based on the nature of the TLR (4nt-, 5nt- and IL-TLR). Since each FMN riboswitch has two TL/TLR interactions, these sequences were divided into two separate alignments, one for L2-L6 and one for L3-L5. Further trimming of these sequences yielded alignments restricted to the TL/TLR interaction and two nucleotides on the 3′ -side of the T-loop-bearing helix that interacts with the TLR (**Supplementary Data**). These alignments are the basis of comparison of sequence space from the selection and that in biology.

This analysis reveals several features of the TL/TLR interaction. First, a breakdown of the sequence identities of each position reveals clear patterns of nucleotide frequencies at each position and consensuses (**Tables S1-S3**). In prior work, this analysis was limited to T-loops observed in crystal structures, providing a limited understanding for informing these patterns.^15^ Within the T-loop motif, we find strong agreement with the consensus derived from structures of 11 introns and riboswitches.^15^ However, the expanded group of sequences within our alignments differ in several important ways. First, with respect to the second nucleotide position, the strict requirement of uridine or guanosine suggested by prior analysis is not supported in all cases. While T-loops interacting with an IL-TLR in class-II Cbl exclusively have a guanosine at that position (**Table S3**), we observe that guanosine is preferred and cytidine and uridine are tolerated in FMN riboswitches and class-I Cbl riboswitches. It is notable that the TL/5nt-TLR interaction, nucleotide position 2 is almost exclusively guanosine. This indicates that sequence variation within the T-loop is dependent on its structural context.

**Table 1:**
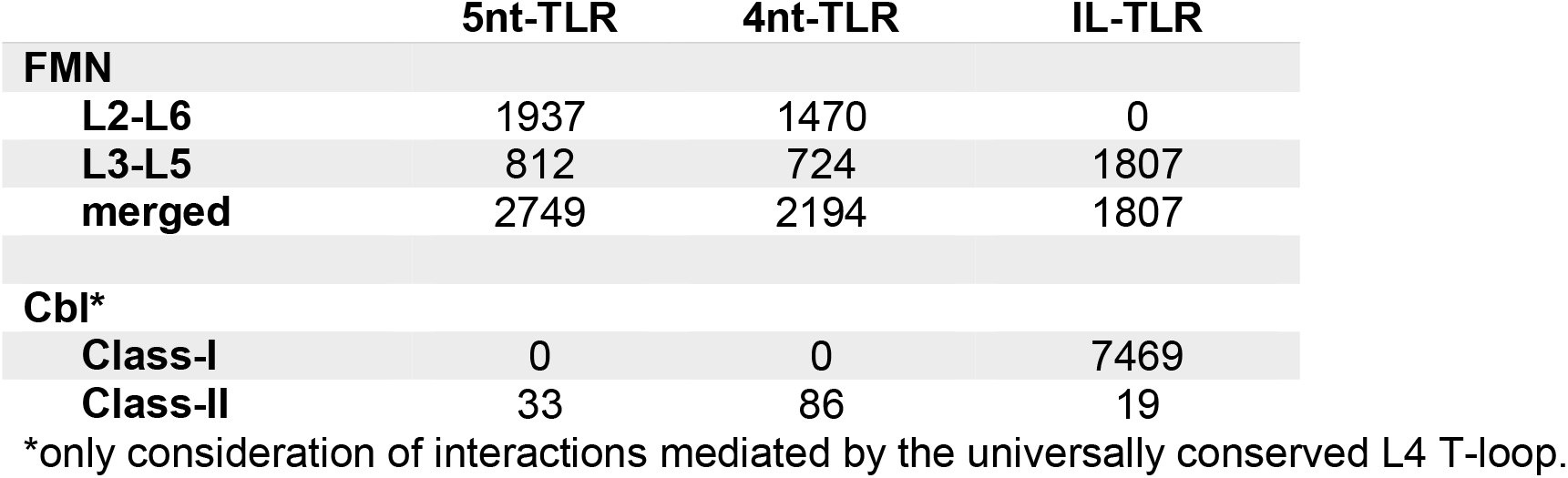
Occurrence of three types of T-loop receptors in FMN and Cbl riboswitches

|  | 5nt-TLR | 4nt-TLR | IL-TLR |
| --- | --- | --- | --- |
| **FMN** |  |  |  |
| **L2-L6** | 1937 | 1470 | 0 |
| **L3-L5** | 812 | 724 | 1807 |
| **merged** | 2749 | 2194 | 1807 |
| **Cbl\*** |  |  |  |
| **Class-I** | 0 | 0 | 7469 |
| **Class-II** | 33 | 86 | 19 |

\*only consideration of interactions mediated by the universally conserved L4 T-loop.

A second aspect of the FMN/Cbl TL/TLRs revealed by this analysis is that the different TLRs are not functionally identical (**Table 1**). For example, within the FMN riboswitches, the L3-L5 interaction uses all three types of TLRs to form the tertiary interaction with a preference for the IL-TLR, whereas the IL-TLR is never observed in the L2-L6 interaction. This would not be due to the additional helix from the IL-TLR sterically interfering with the rest of the structure or with FMN binding, as it is on the surface (**Figure 1(D)**). It is therefore reasonable to conclude that its exclusion is likely due to an aspect of the RNA folding, such as a slower rate of folding that would be deleterious to function. Within class-II Cbl riboswitches, where the three TLRs are observed, there appears to be a significant bias towards the 4nt-TLR, which again may reflect the rate of folding of the RNA rather than structural stability or ligand binding.

### A cell-based screen for sequences supporting functional TL/TLR interactions

We then used a cell-based screen to interrogate the TL/TLR interaction and understand the relationship between its sequence space and function. The basis of this screen is an *E. coli* reporter vector in which the *env*8 Cbl riboswitch is dropped into the 5′ -leader sequence of an mRNA encoding a fluorescent protein (FP) that serves as a readout for regulatory activity.^19^ The *env*8 riboswitch is an OFF switch in which Cbl promotes a long-range tertiary interaction between the aptamer and the ribosome binding site (RBS) that prevents translation. Therefore, bacteria grown in a chemically defined medium will show high levels of fluorescence in the absence of cyanocobalamin (CNCbl) and low levels of fluorescence when the medium is supplemented with CNCbl. To screen for functional TL/TLR sequences, a region of the riboswitch was fully randomized. The number of fully randomized positions was limited to seven to ensure that less than 100,000 colonies need to be screened to guarantee >90% coverage of the library.^69^ The activity of each sequence variant that emerged from the screen was assessed in a liquid culture assay to determine FP expression levels in the presence and absence of CNCbl, which was used to calculate CNCbl dependent activity (raw and processed data collected in this work is given in **Supplemental Data**). This approach has been used previously to investigate another functional T-loop in *env*8.^59^ It is important to note that in this genetic screen, all TL/TLR variants within a given library are hosted in the same riboswitch sequence and thus their relative activities are not confounded by changes elsewhere in the RNA.

For this work, there are two activities that will be assessed to establish function. First, the overall repressive activity in the presence of CNCbl relative to *env*8 is a direct measure of the riboswitch’s ability to fold, bind CNCbl, and occlude the RBS within a limited kinetic window.^60^ Those sequences with lower expression levels of FP in the presence of CNCbl are better performers and for the purposes of analyses in this study, sequences with a relative repression value of <3.0 (i.e., up to 3-fold above *env*8 levels of Cbl-dependent FP repression) are considered strong performers. Because some sequences in the library may repress expression by a Cbl-independent mechanism, the second consideration is the overall fold repression relative to *env*8, calculated as the ratio of the relative fluorescence in the absence and presence of CNCbl. Higher expression ratios relative to *env*8 represent better Cbl-dependent regulatory activity and in this study any variant that displays an expression ratio of >0.5 is considered a strong performer. Thus, sequence variants that strongly support Cbl-dependent regulatory activity will be reflected in both metrics.

### Screening of a library consisting of the T-loop and directly interacting nucleotides

As a first step towards a detailed analysis of the FMN/Cbl TL/TLR interaction, we performed an analysis of the well-characterized T-loop motif and its interaction with nucleobases in its receptor. A library was designed in which the T-loop module corresponding to the first five nucleotides of L4 and two directly interacting nucleotides in L6 were fully randomized (red and cyan, respectively; **Figure 2(A)**). The T-loop (nucleotides 29-33, *env*8) in L4 (red, **Figure 2(B)**) directly interacts with two nucleotides within a seven-nucleotide terminal loop TLR: U60 and A61 (cyan, **Figure 2(B)**). While A61 forms a canonical intercalative interaction with the T-loop along with two hydrogen bonds with the sugar edge of G30, U60 only forms van der Waals contacts with A31 (**Figure 2(C)**).

**Figure 2.**
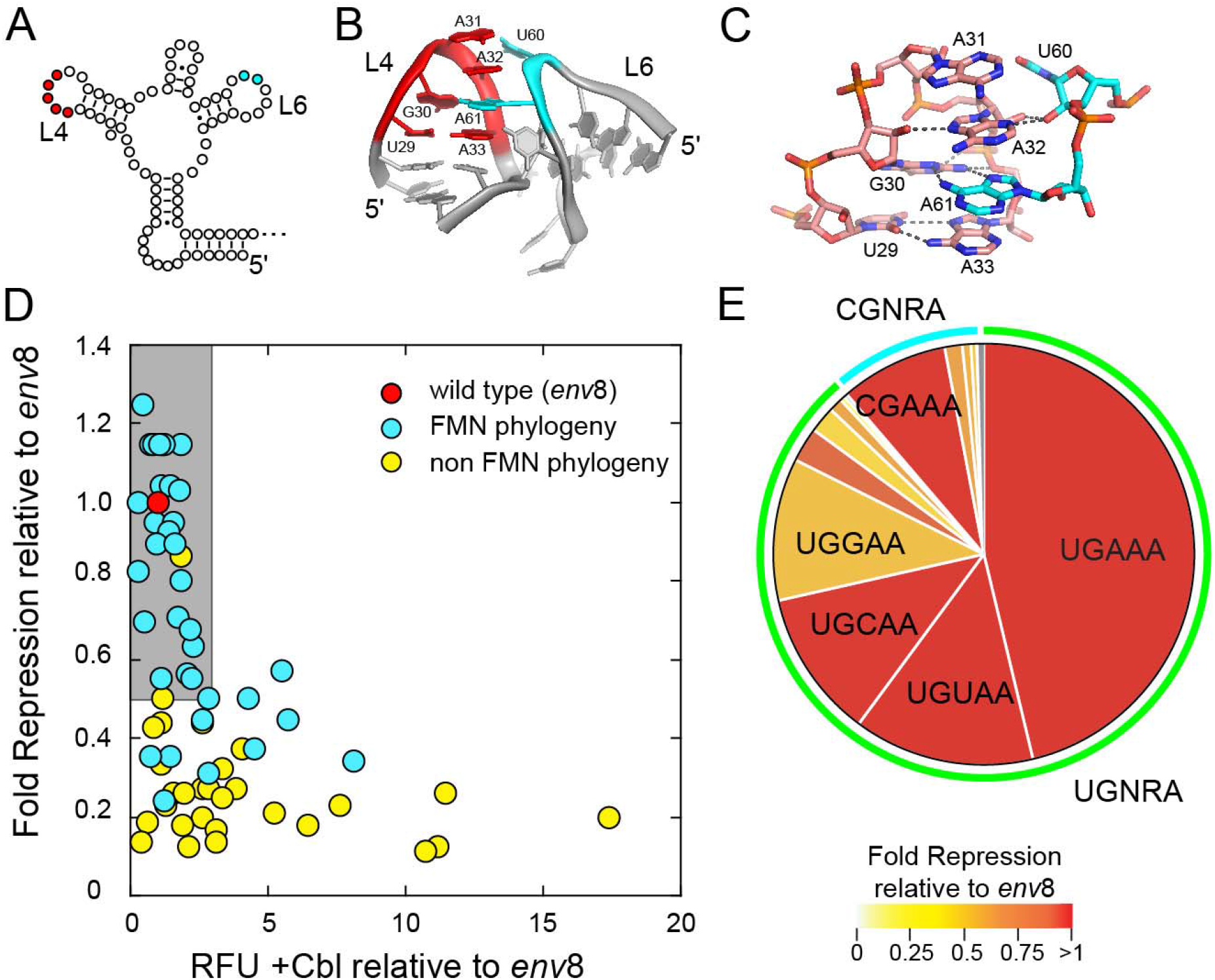
Genetic screen of the *env*8 T-loop and associated nucleotides. (A) Secondary structure of *env*8 aptamer domain with nucleotide positions represented as open circles and the fully randomized regions in the library are colored as in Figure 1. (B) Cartoon representation closeup of the TL/5nt-TLR interaction emphasizing the architecture of the randomized nucleotides (PDB 4FRG). (C) Same perspective as in panel B but showing the nucleobase-mediated hydrogen bonding network between the T-loop and the 5nt-TLR for the nucleotides randomized in this library. (D) Plot of the performance of the sequence variants that emerged from the screen. Individual variants are colored red (*env*8, considered as wild type against which all other variants are normalized), sequence variants found in the sequences of TL/5nt-TLRs in the FMN riboswitch family are cyan, and sequence variants for those not observed in FMN sequence space are colored yellow. Gray box marks those variants with strong performance (i.e., relative fluorescent units (RFU) <3.0 relative to *env*8 and Fold Repression >0.5 relative to *env*8). (E). Pie chart of the core five nucleotide T-loops found in the FMN family that interact with 5nt-TLRs. Each slice represents the fraction of the population that comprises a specific sequence and the color of each slice represents the population-weighted average of the fold repression activity normalized to *env*8 for observed variants in that have that specific T-loop sequence. Gray slices represent sequence space not observed in the selection. Curves around pie chart denote overarching sequence motifs. Green and cyan marks denote TGNRA and CGNRA motifs, respectively.

These seven positions were fully randomized in the screening library and ∼53,300 colonies were screened corresponding to a 96% probability of observing any single sequence within the library (“TL”, **Table S4**). From this screen, 57 unique variants were observed, and their Cbl-dependent activity measured in liquid culture, resulting in 45 variants with at least 10% of wild type activity (sequences of variants from screens performed in this study are provided in a fasta file format; **Supplemental Data**). The Cbl-dependent repressive activity relative to *env*8 versus Cbl-dependent repression relative to *env8* of each of the 45 variants was plotted to visualize the best performing variants (gray box, **Figure 2(D)**). The data points were also colored according to whether the variant sequence was observed in an alignment of 2,749 instances of a T-loop interacting with a 5nt-TLR in the FMN riboswitch family. Expectedly, the best performing variants were almost all represented in biological sequence space and most of the poorly performing variants were not. The two variants that show strong performance that were not observed in the FMN riboswitches, however, are closely related to sequences found within the FMN family and therefore do not represent a different solution to establishing this tertiary interaction. Thus, the cell-based screen correlates well with the observed biological FMN/Cbl T-loop sequence space.

To further assess which specific sequences promote efficient regulatory activity, we examined individual features of this library. First, we considered the T-loop motif, comprising the first five nucleotides of the seven-nucleotide terminal loop. To do this, we determined the relative populations of the different T-loop sequences within the FMN TL/5nt-TLR alignment (**Table S1**) and visualized these data in a pie chart to show the frequency of each observed T-loop (**Figure 2(E)**). Most T-loop sequences fall into two motifs: UGNRA (88.7%) and CGNRA (10.9%) (where N is any nucleotide and R is a purine). All sequences outside these two motifs were singletons (cyan and green rings **Figure 2(E)**). Thus, the FMN/Cbl T-loops have the sequence consensus of YGNRA (where Y is a pyrimidine), although the majority (∼80%) fall into the consensus UGNAA. This consensus excludes other T-loops found in biology such as that of tRNAs that have different sequence features, such as a uridine at the second position rather than a guanosine.^15^ Coloring the population chart according to its observed Cbl-dependent fold repression relative to *env*8 shows that all sequences represented by the YGNRA T-loop motif space support robust regulatory activity (phylogenetic population data used to compose this and other pie charts provided in **Supplementary Data**). In consideration of function, it should be noted that guanosine at position 3 is slightly disfavored, as seen with UGGAA motif, suggesting optimal T-loop sequences have the consensus UGHAA (where H is not G) motif.

The second component of the library that was assessed were the two nucleotides in the 5nt-TLR interacting with the T-loop. The first randomized position, corresponding to U60 (contact *i*, **Figure 1(E)**), displays significant variation across both phylogeny and the genetic screen. Within the dominant T-loop sequence (UGAAA), position 61 is observed in biology as all four nucleotides, with a strong preference for uridine (**Figure S1(A)**). However, all four sequence variants have strong repressive activity (note that uridine at this position corresponds to wild type *env*8), indicating that there is no preference for base pairing between nucleotide positions 31 and 60 despite their spatial proximity. The UGCAA and UGUAA T-loops show a similar lack of preference for 31-60 base pairing. The UGGAA T-loop shows lower activity than the other loops with a functional preference for cytidine, suggesting that this loop may support a G-C base pair (**Figure S1 (A)**). Further, the UGGAA T-loop displays lower activity for two guanosine nucleotides comprising contact *i*, which could be indicative of steric clashing. While the UGNRA T-loops show little preference for the nucleotide at position 60, the CGNRA tetraloops show a very strong bias towards pyrimidines at this position across FMN riboswitches (**Figure S1 (B)**). The prevalent CGAAA T-loop (**Figure 2(E)**) strongly prefers pyrimidines at position 60, although purines at this position also support a near wild type repressive activity. The CGGAA T-loop is observed only in conjunction with pyrimidine nucleotides at position 60, reinforcing this bias. Notably, the most prevalent R-Y combinations are A-U and G-C, suggesting that in the context of the CGNRA T-loops, there is a preference for a Watson-Crick (WC) base pair between the T-loop and TLR. Together, these data suggest an interdependence between position one to five pairing and the nature of contact *i*.

The second randomized nucleotide in the 5nt-TLR, corresponding to A61 (contact *ii*, **Figure 1(E)**), is almost universally conserved in the FMN riboswitches as an adenosine (99.96% A, 0.04% T; **Table S1**) and almost all highly active variants have an adenosine at this position. The only sequence variant observed in this screen with modest activity that does not have an adenosine at this position is a UUAAA T-loop (tRNA-like T-loop) interacting with a cytidine at position 61. Thus, the genetic screen supports biology’s almost exclusive use of adenosine to intercalate between the fourth and fifth positions of the T-loop.

Together, these data strongly support the FMN/Cbl group of T-loops interacting with a 5nt-TLR as having the consensus YGNRA motif, with the sequence UGHAA functionally favored. An alternative projection of the data in which the variants are colored according to consensus sequence reinforces that the best performing T-loops all fall within the YGNRA sequence motif (**Figure S2**). However, we find that not all active T-loops fit within this consensus. For example, we find several sequences that fall into the tRNA T-loop consensus sequence (UUCRA) or a unique rRNA T-loop (GGAAG).^15^ Thus, other groupings of T-loops can support folding and activity of the Cbl riboswitch, though they are classified as weaker performers (yellow, **Figure S2**). The second finding of this work is that while the UGNRA and CGNRA motifs are structurally and functionally similar, they do exhibit differences in their preferences of interacting nucleotides. Specifically, we observe that the UGNRA motif has little sequence preference for nucleotides involved in contact *i*, but that CGNRA motifs show a strong preference for pyrimidines.

### The FMN/Cbl 5nt-TLR motif

To assess the sequence requirements for the 5nt-TLR module, we created a library in which the first five nucleotides of the 5nt-TLR module were fully randomized along with two nucleotides at the 3′ -end of the T-loop that were not assessed in the previous library (cyan and magenta, respectively; **Figure 3(A)**). The rationale for this library design is based upon a similar organization of the 5nt T-loop bearing L4 and the 5nt-TLR bearing terminal loop L6 (**Figure 1(B)**). Like L4, the first five nucleotides of L6 form a structure in which positions 1 and 5 form a non-canonical base pair between U59 and A63, in the same fashion as the T-loop. However, unlike the T-loop, the central three nucleotides (60-62) are splayed out from the loop and have different roles. Nucleotides 61 and 62 (positions 3 and 4 in the 5nt-TLR) are engaged in direct contacts with the T-loop, as described above; randomization of these two nucleotides yielded no nucleotide preference for position 60 and strong preference for adenosine at position 61 (see previous section). The nucleotide at position 60 (position 2) is flipped out towards solvent and does not engage in any direct contacts with the rest of the RNA. Finally, like T-loop bearing L4, the last two nucleotides of L6 are bulged from the rest of the loop to engage in long-range contacts and are therefore distinct from the receptor motif (**Figure 1(B)**). In L4, nucleotide 35 (position 6 of the terminal loop) forms a hydrogen bond contact with the backbone of the TLR and is stacked on the non-canonical U-A pair, while nucleotide 36 (position 7) is flipped out towards solvent and makes no productive interactions with the rest of the RNA (**Figure 3(C)**). Thus, this library comprises the TL-like TLR loop and two nucleotides from L4 that mediate contact *iii* (**Figure 1(E)**).

**Figure 3.**
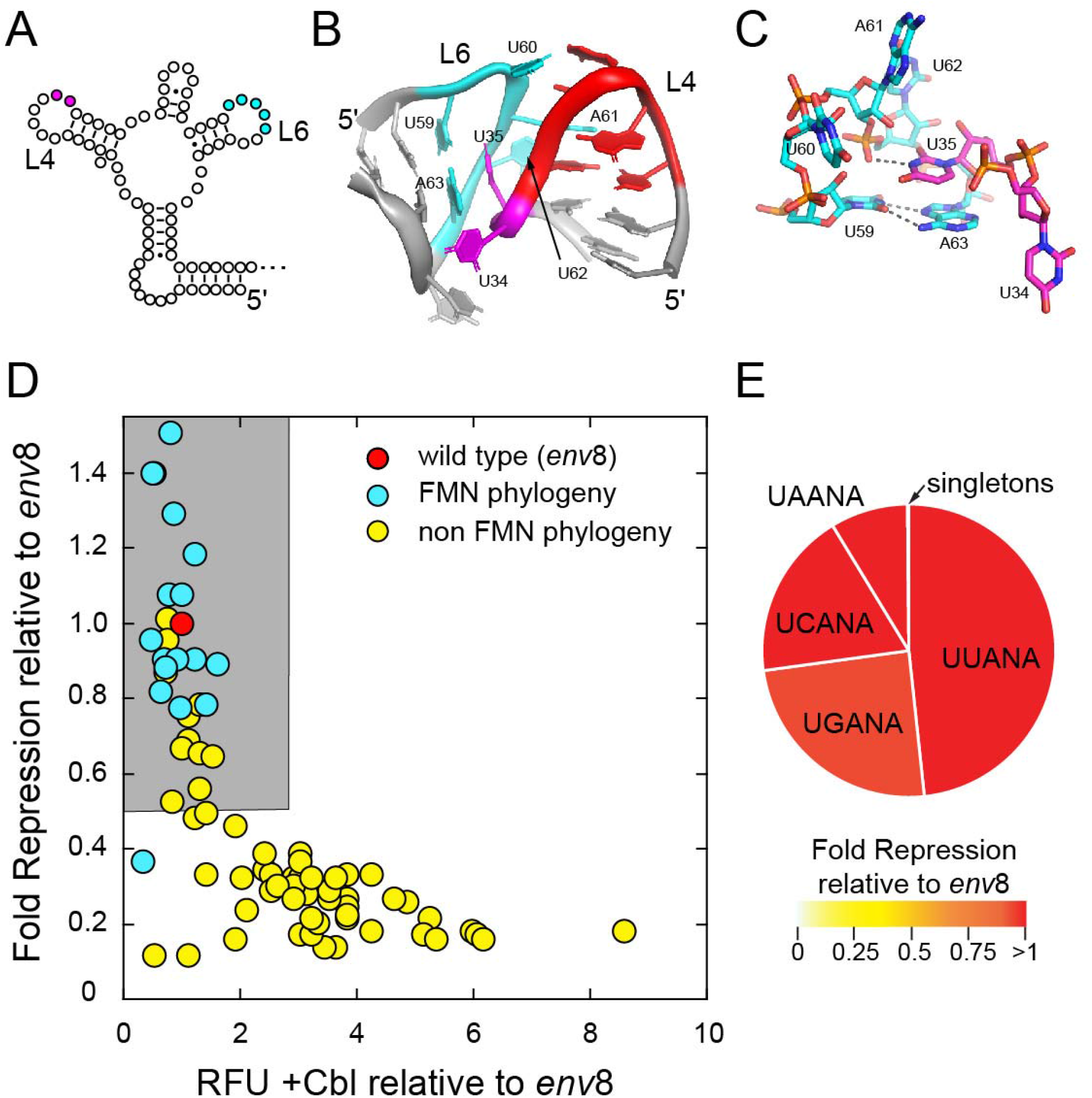
Genetic screen of the 5nt-TLR. (A) Secondary structure of *env*8 aptamer domain with nucleotide positions represented as open circles and the fully randomized regions in the library are colored as in Figure 1. (B) Cartoon representation closeup of the Cbl TL/5nt-TLR interaction emphasizing the architecture of the randomized nucleotides (PDB 4FRG). The numbering of nucleotides is consistent with that of the L4-L6 interaction in the *env*8 riboswitch. (C) Stick representation of the nucleobase-mediated hydrogen bonding network between the T-loop and the 5nt-TLR for the nucleotides randomized in this library. (D) Activity plot of the variants observed in the genetic screen. (E) Pie chart of the population distribution of the core five nucleotide sequence motif of the FMN 5nt-TLR. In panels D and E, the color scheme is the same as in Figure 2.

These seven positions were fully randomized in a separate screening library and ∼49,700 individual colonies were screened, corresponding to 95% probability of observing any one individual sequence. From this screen, 75 unique sequences were identified with 59 displaying Cbl-dependent FP expression (“5nt-TLR”, **Table S4**). As above, the regulatory activity of each sequence was determined and plotted as reporter expression level in the presence of Cbl versus Cbl-dependent fold repression (**Figure 3(D)**).

The dominant sequence motif of highly functional 5nt-TLRs is UNANA, which also comprises almost all sequences observed across FMN riboswitches. The functionality plot reveals two distinct clusters of activity, with a well-populated set displaying both strong repression in the presence of Cbl (<3-fold above *env*8) and strong Cbl-dependent repression (>0.5-fold above *env*8) (gray shaded region, **Figure 3(D)**). This region includes sequences observed in FMN riboswitches as well as a significant number that do not. The only sequence in the FMN family outside of this region is a UUAAA variant. The lower activity of this RNA is due to the L4 interacting nucleotides (CU) and constitutes a solution that yields only reasonable levels of Cbl-dependent regulation but very strongly represses expression. Variants such as this could be needed for genes in which leaky expression of the transcriptional unit being regulated is deleterious to the cell. This highlights how phylogenetic analysis coupled with functional screening can yield variants with divergent properties that are desirable in specific genetic contexts.

An alternative visualization of these data in which each point is denoted by a consensus sequence (UNANA, CNANA, ANANA or others) indicates that the highly active region is almost entirely populated by UNANA sequences (**Figure S3**). There are a few of the CNANA class, underscoring that like T-loops, a non-canonical C-A pair supports function almost as well as the non-canonical U-A pair. However, in FMN phylogeny, the 5nt-TLR is almost entirely represented by the UNANA group, all of which are highly active (**Figure 3(E)**), indicating that this represents the biologically preferred consensus motif. Comparison of the two visualizations reveals that the non-biological sequences within the highly active region fall within a larger consensus of YNANA for the 5nt-TLR, further identifying this as the consensus motif. Finally, these data indicate that the 5nt-TLR shows little preference for the identity of nucleotides at either position 2 or 4. This is consistent with the previous screen which showed little preference at position 2 and the crystal structure that reveals a lack of engagement with other elements of the RNA by the nucleotide at position 4. Across phylogeny, there are biases against some nucleotides such as A at position 2 and G at position 4 in the FMN phylogeny suggesting these nucleotides may impede folding or function to a small extent (**Table S1**).

The two bulged nucleotides of the 3′ -side of the T-loop also exhibit patterns of conservation in the selection that are consistent with their roles in the TL/TLR interaction. In variants arising from the screen, the nucleotide at position 6 of the TL (nucleotide 34 in *env*8) displays a high degree of variation, with all four nucleotides represented (2 A, 6 C, 7 G, and 13 U) within the highly active group of variants. This reflects the solvent exposed positioning of the nucleobase at this position (**Figure 3(C)**). In contrast, the nucleotide at position 7 (nucleotide 35) shows a strong bias towards uridine and purines are not tolerated (1 A, 7 C and 20 U). The insertion of the position 35 nucleotide by the T-loop into the core of the 5nt-TLR is analogous to the donation of an adenosine by the TLR to the T-loop but shows more sequence tolerance. It should be noted that these two positions show more conservation in class-II Cbl and FMN phylogenies. In FMN riboswitches, position 6 is highly variable but is biased against adenosine and towards uridine, and position 7 is almost exclusively uridine (98.5%) (**Table S1**). Thus, from our experimental and biological sequence analysis, the functional consensus for the full FMN/Cbl T-loop bearing terminal loop is (YGNRA)NY where the sequence in parenthesis denotes the T-loop with a more conservative consensus encompassing the most functional sequence space being (UGHAA)UU based on the TL and 5nt-TLR libraries surveyed.

### The FMN/Cbl 4nt-TLR motif

The four nucleotide TLR motif is slightly less represented across FMN phylogeny and is also found in the class-II Cbl riboswitches (**Table 1**). From a structural perspective, the key difference between the two TLR motifs is that the nucleotide equivalent to U60 in *env*8 (position 2 of the 5nt-TLR; dashed circle, **Figure 4(A)**) is deleted such that there is no nucleotide available for contacting the position 3 nucleotide of the T-loop (**Figure 4(B)**). All other features of the TL/TLR interaction are maintained, including donation of a nucleotide to the other loop (i.e., A90 of the FMN TLR and U24 of the T-loop, **Figure 4(B,C)**) as well as the maintenance of a non-canonical base pair between the first and last positions of the motif.

**Figure 4.**
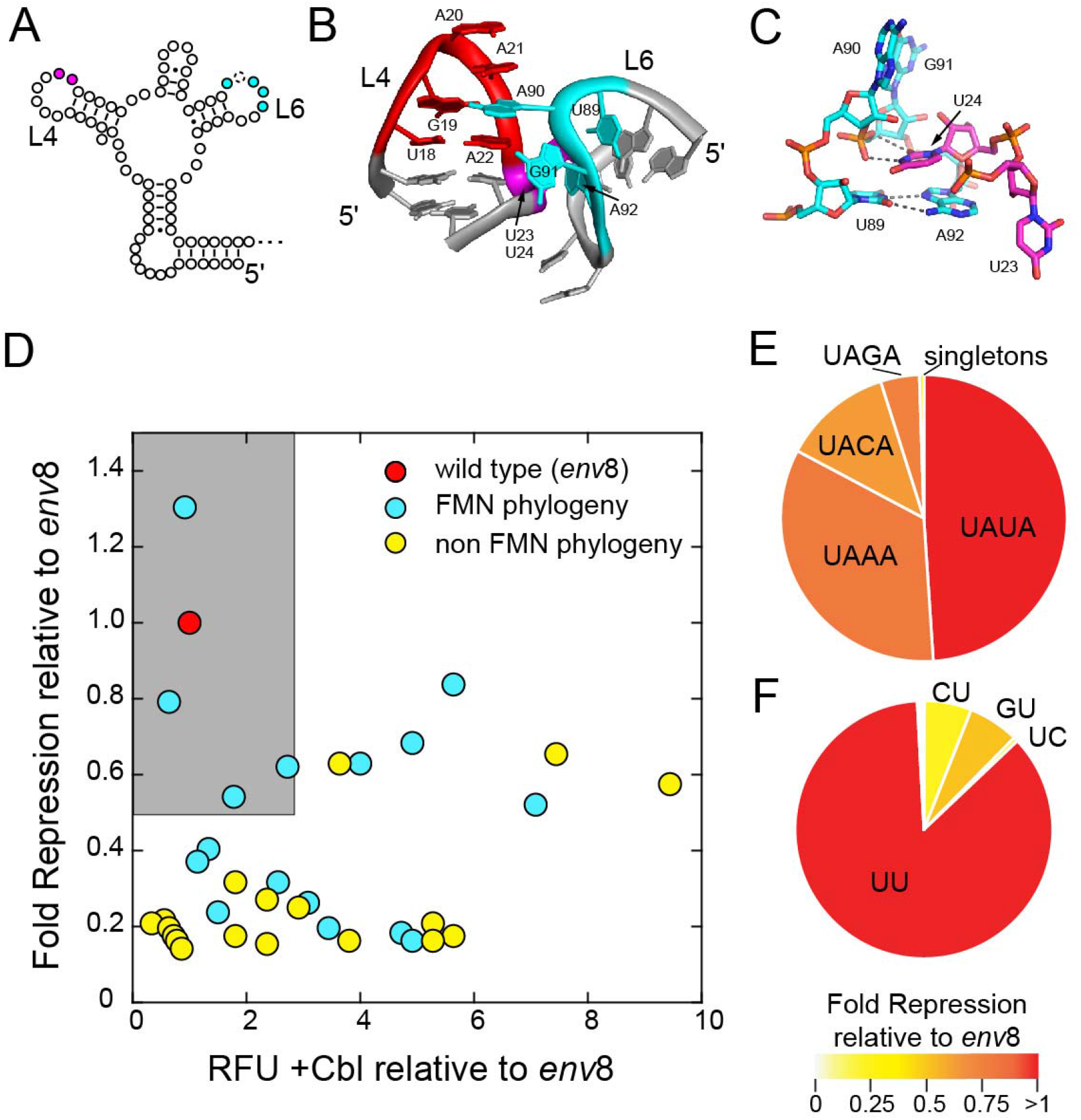
Genetic screen of the 4nt-TLR. (A) Secondary structure of *env*8 aptamer domain with nucleotide positions represented as open circles and the fully randomized regions in the library are colored as in Figure 1. The circle with the dashed line represents the nucleotide deleted in the library. (B) Cartoon representation closeup of the TL-4ntTLR interaction emphasizing the architecture of the randomized nucleotides. (PDB 3F4E) The numbering of nucleotides is consistent with that of the L2-L6 interaction in the FMN riboswitch. (C) Stick representation of the nucleobase-mediated hydrogen bonding network between the T-loop and the 4nt-TLR for the nucleotides randomized in this library. (D) Activity plot of the variants observed in the genetic screen. (E) Pie chart of the population distribution of the core five nucleotide sequence motif of the FMN 4nt-TLR. (F) Pie chart of two nucleotide sequence motif of the FMN T-loop that interacts with the FMN 4nt-TLR. In panels D-F, the color scheme is the same as in Figure 2.

To assess the functional sequence space of the TL/4nt-TLR interaction, we created a library similar to the 5nt-TLR (cyan and magenta positions fully randomized; **Figure 4(A)**). For the resultant screening library (“4nt-TLR”; **Table S4**) ∼18,200 individual colonies were screened, corresponding to 99% probability of observing any one individual sequence. From this screen, 36 unique sequences were identified with 21 displaying Cbl-dependent FP expression. As above, the regulatory activity of each sequence was determined and plotted as reporter expression level in the presence of Cbl versus Cbl-dependent expression (**Figure 4(D)**).

In contrast to the 5nt-TLR library, the functional plot only contains a few sequences that are highly active (gray box, **Figure 4(D)**). These sequences fall within the consensus U-ANA (the dash represents the deleted nucleotide position), which is consistent with the top performers of the 5nt-TLR sequence and the structure of the 4nt-TLR. The U-ANA motif also dominates the FMN phylogeny and represents 99.5% of sequences with the rest of the sequence space mostly represented by singletons (**Figure 4(E)**). The dominant sequence (U-AUA) exhibits performance exceeding *env*8 5nt-TLR, but the others show reduced activity. In contrast to the 5nt-TLR, there are multiple sequences that have strong Cbl-dependent repressive activity (>0.5-fold of *env*8), but weakly repress gene expression in presence of Cbl (**Figure 4(D)**). That is, a number of switches have ∼4-8-fold repression of gene expression between ± Cbl, but they fail to strongly turn expression off and thus could be considered “leaky”. This region of the plot contains sequence variants that are both represented and not represented across FMN riboswitches. Interestingly, all of these “leaky” sequences fall within the U-ANA motif with the exception of a U-UAC 4nt-TLR sequence.

The other randomized component of this library was the two bulged nucleotides on the 3′ -side of the T-loop in P4, as in the 5nt-TLR library (**Figures 1(B) and 4(A)**). Examination of the sequence frequencies at these two positions in the FMN riboswitch family reveals a similar bias as observed in the T-loops that interact with 5nt-TLRs (**Table S2**), and the sequence UU dominates biological sequence space (**Figure 4(F)**). While the UU sequence supports performance comparable to *env*8, other sequences show lower performance. In particular, a cytidine at position 6 (the first of the bulged nucleotides), which occurs with 7.1% frequency across FMN phylogeny, performs poorly compared to the UU variants. Since this nucleotide is flipped out into solution and does not interact with the rest of the RNA, this is likely an example of the bias being towards a nucleotide that either does not support alternative conformations and/or has the lowest energetic penalty for destacking.

### The FMN/Cbl IL-TLR motif

The third type of TLR observed in FMN/Cbl riboswitches is an internal loop (IL) motif rather than the terminal loop of the 4nt- and 5nt-TLRs (**Figure 5(A)**). The IL-TLR motif shares two structural similarities to the other TLRs: a non-canonical U-A base pair (i.e., U77-A150; **Figure 5(B,C)**) and donation of a nucleobase to the T-loop (i.e., A79, **Figure 5(B,C)**). However, it is important to note that the adenosine that is donated by the IL-TLR presents its WC face rather than its Hoogsteen face to the position 2 nucleotide of the T-loop. On the 3′ -side of the IL (proximal to P7) there are two unpaired purine nucleotides that interact with the backbone of the opposite strand (i.e., A78 and G149). Moreover, the two flipped out nucleotides of the T-loop bearing terminal helix (positions 6 and 7) are directed towards the IL-TLR and stacked on each other, in contrast to the previously described terminal loops (**Figure 5(C)**). The position 6 nucleotide of the T-loop (G55) is positioned to make contacts with the minor groove face of the closing base pair of the helix preceding the IL-TLR and the non-canonical U-A base pair. Finally, the position 7 nucleotide of the T-loop (U56) is wedged into the IL-TLR but does not appear to make hydrogen bonds with the receptor (**Figure 5(C)**).

**Figure 5.**
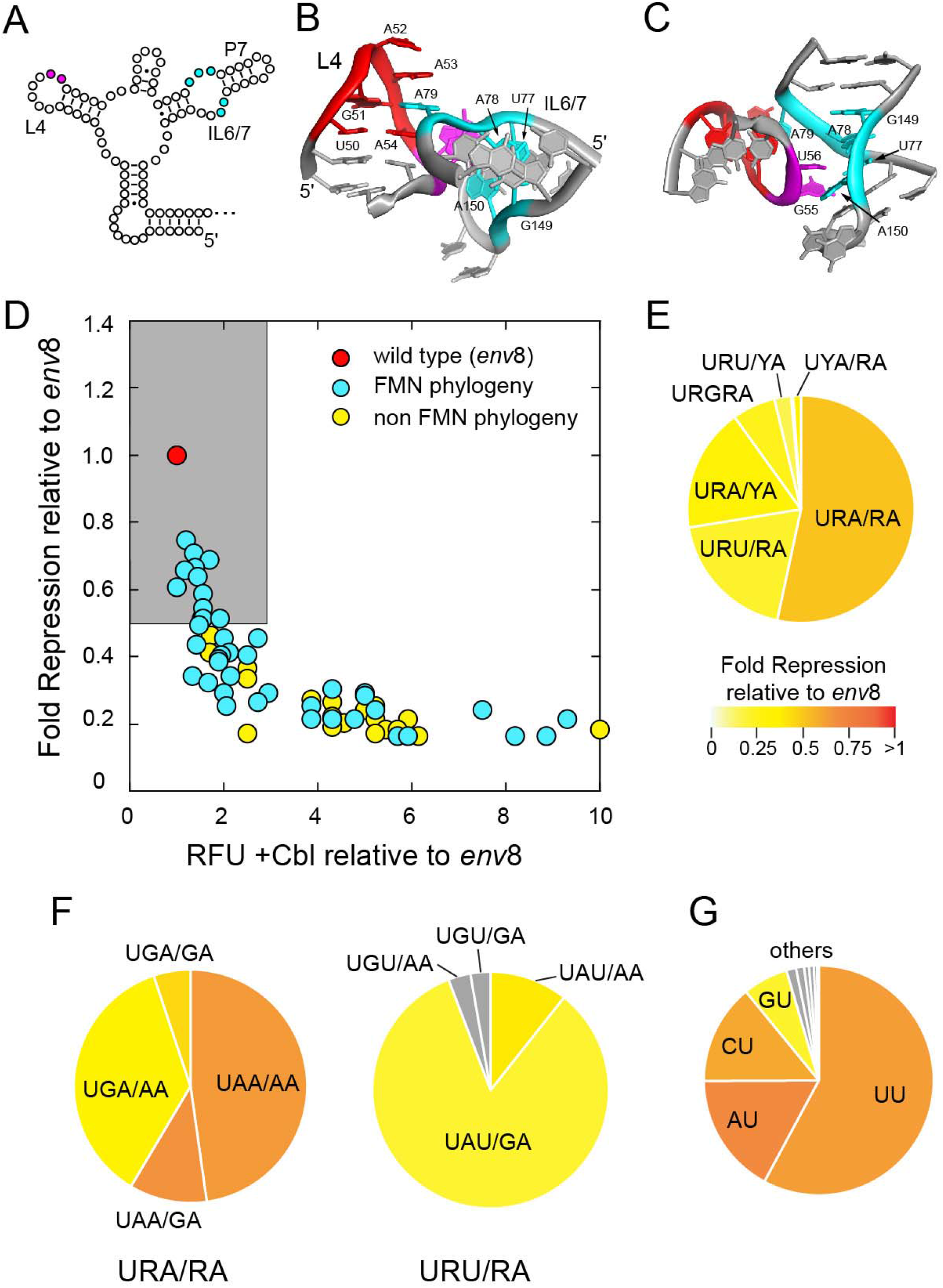
Genetic screen of the IL-TLR. (A) Secondary structure of *env*8 aptamer domain with nucleotide positions represented as open circles and the fully randomized regions in the library are colored as in Figure 1. Note that the secondary structure has changed to reflect the conversion of the L6 terminal loop into an internal loop motif. (B) Cartoon representation closeup of the TL/IL-TLR interaction emphasizing the architecture of the randomized nucleotides. (PDB 4GXY) (C) Cartoon representation closeup of TL/IL-TLR interaction rotated to emphasizing base stacking of randomized area. (D) Activity plot of the variants observed in the genetic screen. (E) Pie chart of the population distribution of the core five nucleotide sequence motif of the FMN IL-TLR. (F) Pie chart of population distribution of FMN riboswitches with the consensus URA/RA (left) and URU/RA (right) with associated activity. (G) Pie chart of the population distribution of sequences of J7/6 for those FMN IL-TLR switches with UAA J6/7 with associated activity. In panels D-G, the color scheme is the same as in Figure 2.

To interrogate the sequence requirements of this interaction in the context of the *env*8 host (which contains a terminal loop TLR), the IL-TLR from the homologous *env*50 class-II Cbl riboswitch was substituted. We have shown previously that this substitution is supported by the Cbl riboswitch with respect to ligand binding and regulatory activity.^62^ A library was generated in which the three nucleotides of the 5′ -strand of the IL (J6/7) and the two 5′ -nucleotides of the 3′ -strand of the IL (J7/6) were fully randomized (cyan, **Figure 5(A)**) as well as the last two nucleotides of the terminal loop hosting the T-loop (magenta, **Figure 5(A)**). Note that the 3′ -nucleotides of the J7/6 strand are bulged out in the same manner as the terminal loop TLRs. For this screening library (“IL”; **Table S4**) ∼67,100 individual colonies were screened, corresponding to 98% probability of observing any one individual sequence. From this screen, 55 unique sequences were identified with 55 displaying Cbl-dependent FP expression. As above, the regulatory activity of each sequence was determined and plotted as reporter expression level in the presence of Cbl versus Cbl-dependent fold repression (**Figure 5(D)**).

Several observations emerge from the analysis of the IL-TLR library. First, all of the IL-TLR variants underperform relative to the terminal loop TLR motifs (**Figure 5(D)**). Nevertheless, they still support Cbl-dependent activity, reflecting their representation in the class-II Cbl riboswitches (**Table 1**). Since the replacement of the terminal loop L6 with an IL6/7-P7 extension may not be fully supported by *env*8, we cannot discount the possibility that the cause of this underperformance is due the lack of full portability of this receptor between different RNAs. In light of this, it is notable that the IL-TLR motif is entirely absent in the FMN L2-L6 interaction (**Table 1**), despite it theoretically being able to be structurally accommodated based upon a lack of steric clash with the rest of the RNA. This suggests that the L2-L6 position may have more stringent requirements for the folding and/or stability of this long-range interaction that can only be conferred by terminal loop TLRs. Nonetheless, we find that the top performers emerging from the selection still perform well by our previously established criteria.

Strikingly, we find biological sequences are distributed throughout the observed range of activities, including a number that perform marginally in the context of *env*8 (**Figure 5(D)**). Within the cluster of top performing IL-TLRs, we observe that the sequences are limited to the URA/RA and URU/RA sequence motifs in J6/7 and J7/6. These motifs dominate the biologically represented IL-TLR sequences, with the URA/RA showing the best performance of the IL-TLR family (**Figures 5(E)** **and S4**). Within these motifs are differences in preferences for the purine nucleotides. In the URA/RA motif, there is a strong preference for an adenosine at position 4, and little preference between guanosine and adenosine at position 2 (left, **Figure 5(F)**). This is surprising in light of crystal structures of this motif, in which the position 2 guanosine forms two productive hydrogen bonds with the opposite backbone; potentially a small repositioning of adenosine may enable a single hydrogen bond. Similarly, the favorable hydrogen bonding interaction between the guanosine at position 4 and the opposite backbone would not be supported by the preferred adenosine. Thus, it appears that the main role of these purines is to maintain the cross-strand purine stack between positions 2 and 4 of the IL-TLR which may be most readily accomplished by adenosines. This may reflect a need for conformational flexibility in the motif to promote docking of the T-loop. In contrast, the URU/RA motif shows a strong preference for an adenosine at position 2 and a guanosine at position 4 (right, **Figure 5(F)**). There is broader sequence variation within a slightly lower performing cluster (**Figure S4**). Unlike the 4nt- and 5nt-TLRs, the observation that the IL-TLR can donate a uridine to the T-loop reflects the different orientation of this nucleobase relative to position 2 of the T-loop. Within the class-II Cbl riboswitches, 17/19 variants fall into the URW/RA sequences with the other two as UAA/UA.

Like the terminal loop TLRs, the IL-TLR interacts with two bulged nucleotides at the 3′ -end of the T-loop containing terminal loop (magenta, **Figure 5** **(B,C)**). Analysis of the conservation patterns of these nucleotides in FMN riboswitches containing the UAA/AA IL-TLR reveals that most sequences prefer a uridine for the second nucleotide, but smaller preference for the first (**Figure 5G**). The activity of the four primary variants show that these riboswitches have good regulatory activity, with the exception of GU, which exhibits more modest activity. The disfavoring of guanosine at position 6 is not clear from the structure, as the role of this nucleotide is stacking upon the uridine in the second position, which should be favored. Instead, disfavoring guanosine may be the result of it promoting an alternative base pairing or structure in the terminal loop containing the T-loop. From the above analysis, the consensus for the IL-TLRs is URA/RA, with the most highly functional variants adhering to the URA/GA motif and interacting with a uridine from the 3′ -end of the T-loop terminal loop.

### Requirement of Watson-Crick pairing between the TLR and nucleotides 3′ of the T-loop hairpin

A universally conserved feature of the TL/TLR interaction found in the FMN/Cbl group is a long-range interaction between two nucleotides at the 3′ -side of the TLR and two single-stranded nucleotides that are immediately 3′ of the T-loop containing stem-loop structure that forms the auxiliary helix (contact *iv*, **Figure 1(E)**). In the wild type *env*8 Cbl riboswitch, nucleotides in J4/5 and L6 (dark blue dots, **Figure 6(A)**) form two WC base pairs that stack beneath the base of the L4 stem (highlight in yellow, **Figure 6(B)**). This interaction is not universal to all T-loop-mediated interactions. For example, in tRNA, only one of the equivalent nucleotides engages with a stacking interactions with an unpaired nucleotide on the 5′ -side of the T-loop hairpin and the other is exposed to solvent.^42^ In the hatchet ribozyme, the TLR does not have any nucleotides that engage in a structurally analogous interaction.^35^ Thus, while this accessory contact that supports T-loop-mediated RNA-RNA interactions appears to be essential for some TL/TLR interactions, contact *iv* is not a universal feature.

**Figure 6.**
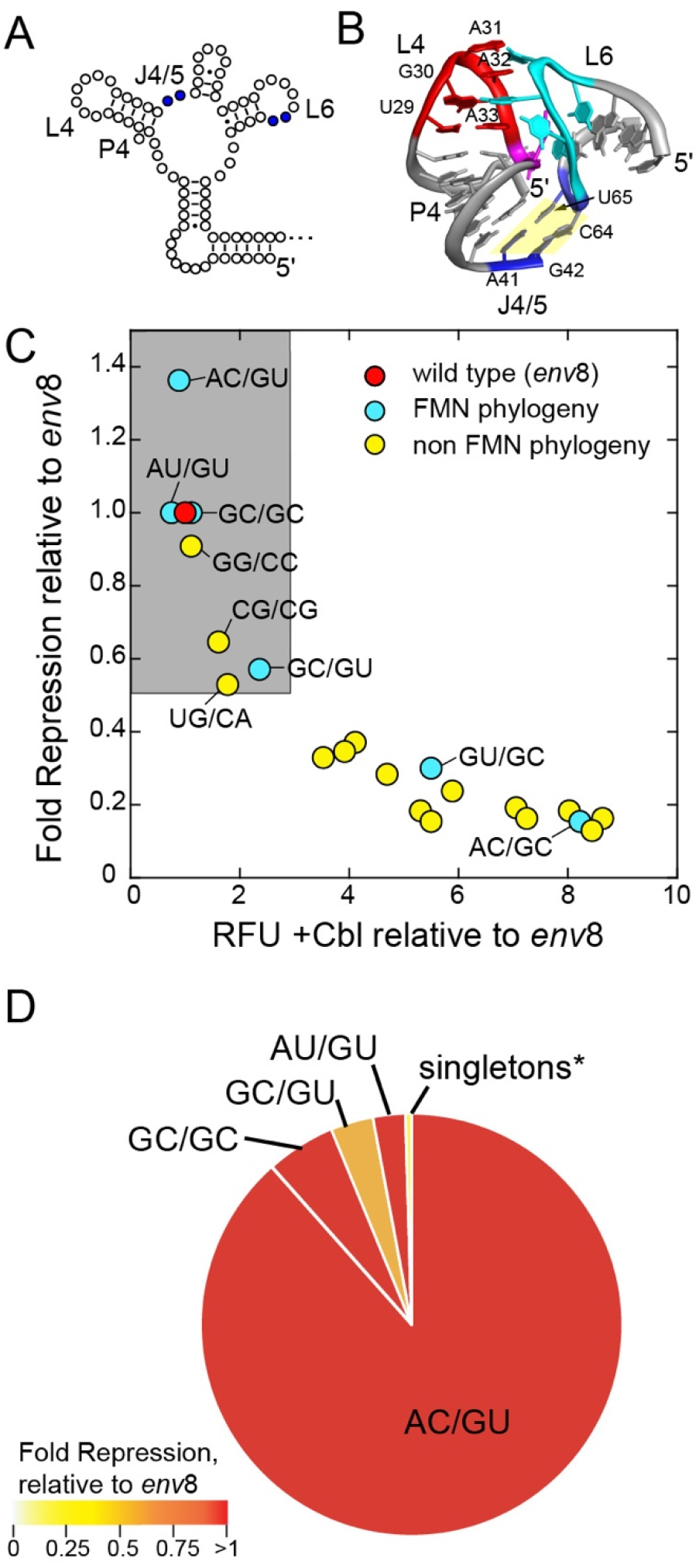
Genetic screen of the auxiliary helix. (A) Secondary structure of *env*8 aptamer domain with nucleotide positions represented as open circles and the fully randomized regions in the library are colored as in Figure 1. (B) Cartoon representation closeup of the TL/5nt-TLR interaction emphasizing the architecture of the randomized nucleotides. (PDB 4FRG) (C) Activity plot of the variants observed in the genetic screen. (D) Pie chart of the population distribution of the auxiliary helix from the FMN TL/5nt-TLR alignment. In panels C and F, the color scheme is the same as in Figure 2.

To assess the importance and patterns of nucleobase-nucleobase interactions required for this interaction in the FMN and Cbl riboswitches, we performed a genetic screen in which the four nucleotides were fully randomized. The resultant library was screened sufficiently so that the probability of observing an individual sequence was >95% (“Aux_helix”, **Table S4**) and 16 unique functional sequences were found. Of these sequences, a cluster with both strong repression in the presence of Cbl and strong Cbl-dependent repression (gray box, **Figure 6(C)**). All of these sequences represented either WC or wobble base pairing between positions 1 and 4 as well as 2 and 3 of the randomized set.

While the screen reveals that WC or wobble base pairing between positions 1 and 4 is favored, the identity of the pair is variable. This likely reflects the fact that this base pair does not interact with other elements of the RNA. However, we do observe a strong preference for R-Y versus Y-R, with all but one of the strong performers having the R-Y configuration (**Figure 6(C)**). In the TLR phylogeny, this preference is absolute. The first base pair is always R-Y with a very strong bias towards A-U in FMN and a smaller bias towards G-C in class-II Cbl riboswitches (**Table S5**). In the other TLR motifs, this preference for A-U versus G-C changes. For example, in the IL-TLR group in FMN, this preference changes from A-U to G-C, while in class-I Cbl riboswitches the IL-TLR group has a nearly equal distribution of A-U and G-C base pairs (**Table S5**). A structural rationale for the R-Y bias is that the nucleotide at position 1 stacks between two nucleotides whereas the position 4 nucleotide only partially stacks on a single nucleotide on its 5′ -side. However, it is not clear why FMN and Cbl riboswitches prefer different base pairs between positions 1 and 4, thought it may reflect folding requirements of the RNAs or subtle structural features.

In the genetic screen, the base pairing between positions 2 and 3 was restricted to either C-G, G-C or U•G pairs. This bias is likely dictated by the fact that an adenosine nucleotide always forms a type-I A-minor triple with the 2-3 base pair.^70, 71^ A thermodynamic analysis of an A-minor triple in the P4-P6 domain of the *T. thermophila* group I intron revealed a strong preference for interactions with C-G base pairs.^72^ The bias towards C-G pairs being the receptor of the adenosine of the type-I A-minor triple is also observed in the *H. marismortii* 50*S* rRNA.^71^ This is strongly supported by phylogenetic analysis, which shows a high representation of C-G base pairs (96.9%) as compared to U-A (0.15%) and U-G (2.5%) wobble pairs (**Figure 6(D)**). However, in the genetic screen, we found other sequences that performed strongly (gray box, **Figure 6(C)**). In particular, we found three sequences having G-C base pairs that are rarely observed in biology, suggesting that this C-G bias cannot be fully rationalized from structural or thermodynamic considerations. Again, given that the activity of riboswitches is typically kinetically constrained, including *env*8,^60, 73, 74^ this bias may be rooted in the folding process. Mapping the repressive activity of the different sequence variants onto a chart showing the relative populations of each biological sequence reinforces the notion that biological sequence space is dominated by the most highly active variants (**Figure 6(D)**). This serves to reinforce the marked preference for the AC/GU sequence motif, although others can support functional almost as well, including the very rare AG/CU motif found in *env*8.

### The top TL/5nt-TLR interactions within the FMN riboswitch family strongly support Cbl-dependent regulatory activity

While our genetic screening approach has the ability to completely examine sequence space in small regions of the RNA, this method precludes examining variants in which nucleotides in separate spatial regions deviate from the *env*8 TL/5nt-TLR sequence. To examine a few of these variants, we first took the alignment of all representatives of the TL/5nt-TLR interaction in the FMN riboswitch family and determined the most represented sequences. These sequences, which represent ∼30% of total sequences (inset, **Figure 7**), strongly conform to the consensus (UGAAA)UU/AC/UGAAAGU, in which individual sequences deviate from that consensus in 1-3 positions. The most conserved aspects of this interaction, such as non-canonical pairs in the T-loop and TLR and the auxiliary helix, are conserved, with variation occurring in regions observed to vary in our screens. However, only variant 5 (v5, inset, **Figure 7**) was represented in the above libraries and was observed to have high activity (top performer in the auxiliary helix library). These sequences therefore represent variation that was not observed in the above screens. As such, we can now address the question of whether these sequences perform better than those where only one component of the interaction was varied relative to the rest of the total sequence of the TL/5nt-TLR interaction.

**Figure 7.**
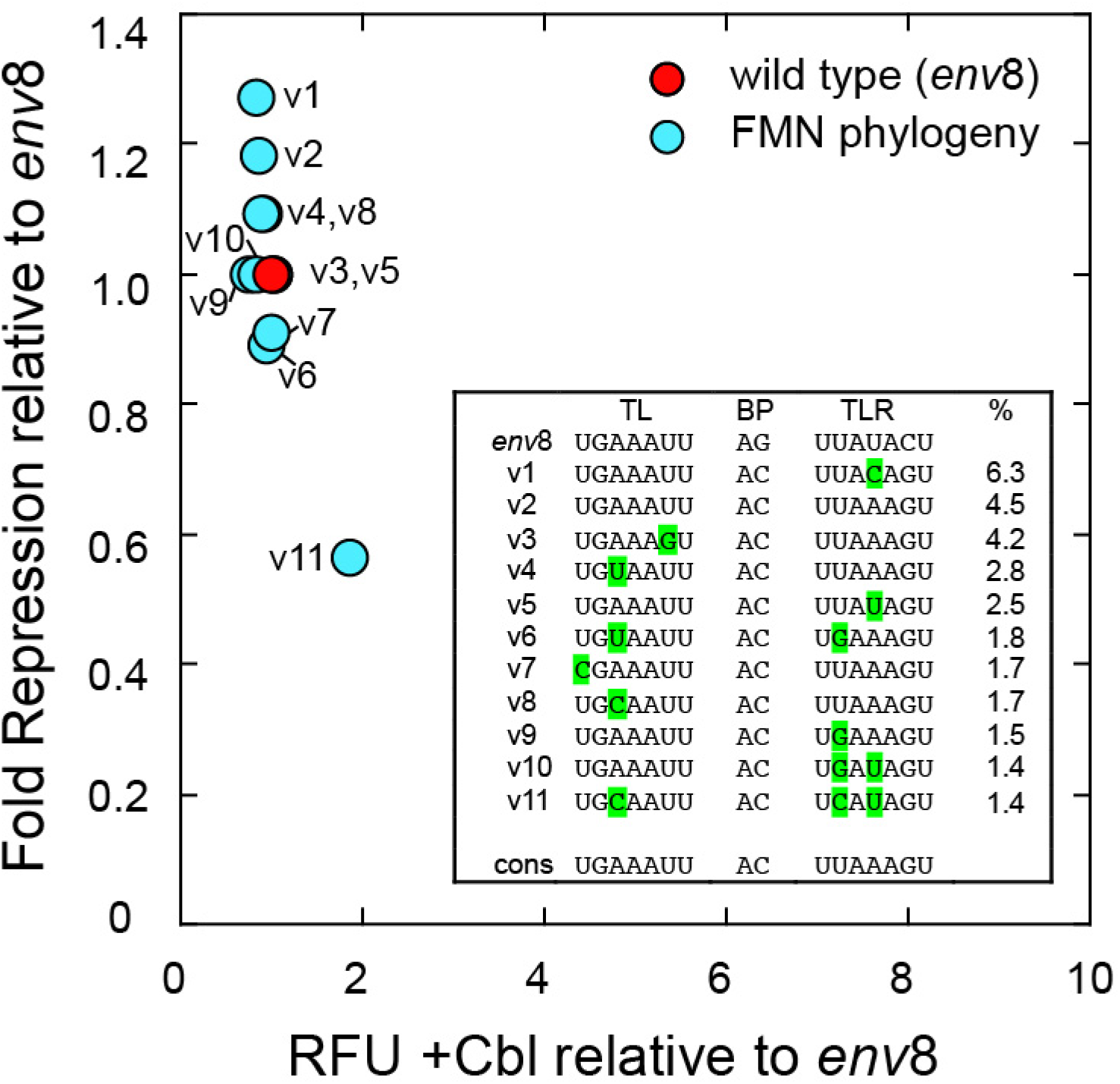
Assay of the common TL/5nt-TLR sequences in the FMN riboswitches. Analysis of the most common sequences in the FMN riboswitch family yielded a set (v1-v11; inset) that comprise ∼30% of all sequences. Nucleotide positions that differ from the wild type *env*8 sequence in the terminal loops are highlighted in green, except positions 2 and 3 in the auxiliary helix which are always a C-G base pair rather than the G-C pair in *env*8. The performance of each variant is plotted as previously, and the labels of the plot correspond to the sequence labels in the inset.

The top sequences in the FMN riboswitch family were synthesized in the context of the *env*8 riboswitch and tested for their ability to support Cbl-dependent FP repression in *E. coli* (**Figure 7**). All of these sequences are considered strong performers by the metrics used for the other libraries, consistent with the findings of the above screening. There are two regions of variation in the top sequences: nucleotide 3 of the T-loop, the spatially proximal nucleotide 2 of the 5nt-TLR, and nucleotide 4 of the 5nt-TLR, which is flipped out into solution. Variation at these positions does not effect riboswitch activity, consistent with the above screens. Most importantly, these data reinforce the boundaries of the sequence space that support high performance TL/TLR interactions in the context of Cbl and FMN riboswitches. These features diverge from the sequences of other T-loops and again suggest that there is a significant context dependence to the T-loop module, imposed by the long-range interactions that it mediates as well as constraints imposed by rapid, high-fidelity folding.

### Conclusions

In this study, we set out to determine the full sequence space that supports the TL/TLR interaction observed in FMN/Cbl riboswitches. Using a cell-based screen, we were able to fully examine sequence space across all nucleotides involved in establishing this critical tertiary interaction. A comparison of the functional sequence space from the screen and sequence preferences in FMN/Cbl riboswitches revealed a set of strong preferences for specific features within the TL/TLR interactions. In particular, there is a strong requirement for a Y-A Watson-Crick/H base pair between positions 1 and 5 of the T-loop in the three TLR motifs. While we find that contact *i* (**Figure 1(E)**) between the T-loop and TLR is not important, as evidenced by its deletion in the 4nt-TLR motif, while contacts *ii*, *iii,* and *iv* (**Figure 1(E)**) have very strong sequence requirements. Notably, but not unexpectedly, the experimentally and bioinformatically observed sequence biases differ between the different TLRs. For example, the 4nt- and 5nt-TLRs almost exclusively donate an adenosine to the T-loop in contact *ii* (**Figure 1(E)**), but the IL-TLR has a relaxed requirement for the donor nucleotide. These observations justify the separation of the TLRs into different groups. Together, these observations suggest that an alternative approach to grouping T-loops by the type of RNA in which they are found^15^ is by the types of contacts that they and their interacting partner engage in—i.e., contacts *i* – *iv*. For example, in the *env*8 riboswitch the T-loop of L4 would be in a grouping of T-loops engaged all four types of contacts while the T-loop in J1/3 would be in another as it only engages in contacts *i* and *ii* via interactions with J6/3.

One aspect of the three groups of TL/TLR interactions that was not interrogated in this study is whether different types of TLRs influence the functional T-loop sequence space. This is because the T-loop used to examine the different TLRs was the same (i.e., UGAAA). However, it is highly likely that sequence space in the T-loop is influenced by the sequence of the interacting TLR. In an analysis of the sequence composition of T-loops found in different contexts, it was observed that there are distinct conservation patterns and nucleotide preferences between the different T-loops. In particular, the T-loops of tRNAs show a distinct pattern of conservation that differs at position 2. Similarly, a cell-based screen of the other T-loop motif in *env*8 (TL1, **Figure 1(D)**), we observe that the identity of position 3 is critically important for regulatory function as it forms a Watson-Crick base pair with a nucleotide in the Cbl binding pocket.^59^ Thus, the T-loop is context dependent and has distinct sequence preferences based upon its interacting partners. In the case of the 5nt-TLR, the sequence requirements at positions 1, 2, and 5 are strong while position 3 is flexible, yielding the observed YGNRA consensus sequence motif.

This context-dependence of subgroups within a motif has practical implications for using these modules to design novel RNAs^11^ such as the recently developed program RNAMake.^14^ This algorithm uses a motif library using junctions, hairpins, and tertiary contacts derived from crystal structures.^14^ Improvements in annotation and classification of these motifs that account for structural context would facilitate the decision making process with respect to incorporating specific modules into a design. In particular, the experimental approach for developing consensus sequences used in this study can help account for dynamic properties of the module that are not apparent from structural snapshots such as folding. Given that kinetic control of riboswitches places strict temporal constraints on their ability to rapidly acquire tertiary structure, functional analysis of motif sequences has the potential to enable the databases to be populated with high performance modules whose ground-state structures can be modeled based upon existing crystal structures. Since riboswitches harbor diverse motifs such as kink-turns,^75^ loop E motifs,^76, 77^ kissing loops,^19, 76^ and multi-helix junctions,^78^ they could be used to functionally annotate the sequence space of these motifs. The above study demonstrates that biology has found highly functional TL/TLRs that confer robust gene regulation and our screened has warranted insights into principles that can allow for informed design of synthetic RNAs.

## Material and Methods

### Library construction through molecular cloning

Riboswitch RNA constructs were amplified through gap-fill PCR using overlapping oligonucleotides synthesized by Integrated DNA technologies. Each library had either 4, 6, or 7 randomized nucleotides and a 5′ NsiI cut site and 3′ HindIII cut site (sequences of all DNA oligonucleotides used in this study are provided in **Table S6**). Constructs of specific riboswitch sequences (e.g., such as the most frequent TL/TLRs in the FMN phylogeny) were created using recursive PCR. Inserts were then digested according to manufacturer’s protocol with Nsi-I HF and Hind-III HF and ligated into a calf-intestinal phosphatase treated plasmid using T4 DNA ligase (New England Biolabs). The resulting recombinant plasmid contained an ampicillin resistance cassette and the riboswitch insert located upstream of the fluorescent reporter protein mNeonGreen.^79^ The plasmid’s transcription is controlled by the moderately strong synthetic promotor ‘<u>proD</u>’^80^ which has previously been used to study the *env8* class-II Cbl riboswitch.^59, 60, 62^ Libraries were transformed into *E. coli* strain BW25113 (Δ *btuR*) cells by adding 3 µL of ligation reaction to a 100 µL aliquot of chemically competent cells, incubated on ice for 15 min, heat shocked at 37 °C for 90 s, incubated on ice for another 5 min, and outgrown in 900 µL of 2xYT rich media for 1 h. Colonies were grown by adding 100 µL of transformant to chemically defined salt broth (CSB) plates that were 1.2% agar supplemented with 100 µg/mL carbenicillin and 5 µM CNCbl. Plates were incubated overnight at 37 °C followed by a 24 h incubation at 4 °C. Photos of colonies were taken under light source with a 490 nm excitation and 510 nm emission filter and used to created counting statistics with the OpenCFU software package (3.9.0).^81^ Percent of potential colonies observed was calculated using an algorithm based on Poisson statistics using the counted colonies observed and the number of possible variants based on the library size.^69^

### Cell-Based Activity Screening

Colonies were transferred to gridded plates and grown overnight at 37 °C. Colonies were illuminated with the same filters as in the initial screen and those that showed differential expression of mNeonGreen on the liganded (dim) and unliganded plate (bright) were selected. Plasmids were isolated from overnight cultures by miniprep (E.Z.N.A® Plasmid DNA Mini Kit I, Omega Bio-Tek) and sequenced by Sanger Sequencing (QuintaraBio).

Those colonies that appeared to display functional switching grided plates were then subjected to activity-based reporter assays. Colonies were inoculated in 3 mL of CSB supplemented with 100 µg/mL of ampicillin overnight at 37 °C. Those outgrowths were then transferred to two secondary 3 mL CSB cultures at a 1:100 dilution, again with 100 µg/mL of ampicillin but one tube was supplemented with 5 µM CNCbl. Secondary cultures were grown at 37 °C until an OD_600_ of 0.4-0.6 was achieved. Activity was determined by measuring cell density (i.e., OD_600_) and fluorescence intensity of 300 µL aliquots of each culture in a Costar® 96-well half area microplate. Plates were read with an excitation wavelength of 490 nm and emission wavelength of 517 nm using a Tecan Infinite M200® PRO plate reader and measured against a maximum fluorescence of 300 µL of a 5 µM fluorescein standard. Each activity assay included single tubes of potential switch colonies as well as an empty vector control (pBR327) and the parental functional switch (*env8*). All measurements were done in technical triplicate, three well per plate per reading, and biological triplicate, three separate measurement days, resulting in at least nine measurements for each riboswitch.

### Data Analysis

Fluorescence values were corrected by the cell density by dividing the fluorescence value by the OD_600_ for each well. Those corrected values were calculated in excel (Microsoft) and then formatted into a csv with cell density corrected values for all colonies detailed in the R markdown for this analysis. Data of all libraries were done in parallel in R to standardize across libraries. Data points that were more that 1.5-times the interquartile range from the upper or lower quartile were removed, a metric chosen assuming a Gaussian distribution.^82^ Following removal of outliers, riboswitches with a fold induction of ≤ 2.0 were removed. Final values were reported as the median repression, expression, and fold induction value and the standard error of the repression and expression values were reported. Finally, FASTA files were created using the values from analysis in R and the unique sequence files for each colony.

### Sequence alignments of FMN and Cbl riboswitches

To construct the alignments used in this study, the full sequence alignments of the FMN (RF0050), Cbl (class-I, RF00174), and AdoCbl-variant (class-II, RF01689) were obtained from Rfam 14.0.^63, 64^ The initial sequence alignments were manually curated to remove sequences that did fully encompass the riboswitch aptamer domain or had deletions in regions that are critically important for structure and/or function. The remainder of the sequences were then re-aligned using MUSCLE.^67, 68^ After multiple iterative rounds of manual curation, the T-loop and TLRs in each alignment were examined and variants that contained insertions or deletions in the motifs were removed, followed by another re-alignment. The class-II Cbl and FMN alignments were then divided into groups based upon the three types of TLRs. Since the FMN riboswitch has two TL/TLR interactions, this sequence was broken into two alignments: one with P3-P5 and another with P1-P2/P6, and each further divided into the three TLR groups. Finally, the sequences in each group were trimmed to contain only the loop motifs and two nucleotides in a joining region that participate in the auxiliary helix. To merge the P3-P5 and P2-P6 TL/TLR files, the P3-P5 sequence had to be circularly permutated to enable P2 to align with P5 (T-loops) and P3 to align with P6 (TLRs). Final alignment files are provided in **Supplementary Data**.

## CRediT authorship contribution statement

**Lisa Hansen:** Methodology, Validation, Formal analysis, Investigation, Data Curation, Writing – Review & Editing. **Otto A. Kletzien:** Investigation, Formal analysis. **Marcus Urquijo:** Investigation, Formal analysis. **Logan T. Schwanz:** Investigation, Formal analysis. **Robert T. Batey:** Conceptualization, Methodology, Validation, Writing – Original Draft, Writing – Review & Editing, Supervision, Project administration.

## Acknowledgements

This work was supported by the National Institutes of Health (Grant no. R01 GM073850 to R.T.B.). The authors would like to thank the members of the Batey laboratory for their discussions and support of this work and particularly to Dr. Lukasz Olenginski for his work on editing this manuscript.

## Appendices

Supplementary Information and Supplementary Data.

