## Supplementary Information for "Context-dependence of T-loop mediated long-range RNA tertiary interactions"

**Lisa N. Hansen, Otto A. Kletzien, Marcus Urquijo, Logan T. Schwanz and Robert T. Batey***

Department of Biochemistry, University of Colorado, Boulder, CO, 80309-0596, USA

**This file includes:**

Supplementary Figures S1 to S4

Supplementary Tables S1 to S6

**Supplementary Material contents:**

**Table S1.** TL/7nt-TLR nucleotide frequencies and consensus.
**Table S2.** TL/6nt-TLR nucleotide frequencies and consensus.
**Table S3.** TL/IL-TLR nucleotide frequencies and consensus.
**Table S4.** Data collection statistics for each library screen.
**Table S5.** Biological variation in the auxiliary helix.
**Table S6.** Sequences of oligonucleotides used in this study.

**Figure S1.** Nucleotide preferences for nucleotides of the 5nt-TLR interacting with the T-loop.
**Figure S2.** Distribution of most prevalent sequence motifs in the T-loop genetic screen.
**Figure S3.** Distribution of most prevalent sequence motifs in the 5nt-TLR genetic screen.
**Figure S4.** Distribution of most prevalent sequence motifs in the IL-TLR genetic screen.

**Table S1. TL/5nt-TLR nucleotide frequencies and consensus**

Cobalamin, class II riboswitch (33 sequences)

| | T-loop | | | | | | | J2/3‡ | | T-loop receptor | | | | | | |
|---|---|---|---|---|---|---|---|---|---|---|---|---|---|---|---|---|
| | 1* | 2 | 3 | 4 | 5 | 6 | 7 | 1 | 2 | 1 | 2 | 3 | 4 | 5 | 6 | 7 |
| A | 91** | 0 | 82 | 100 | 100 | 6 | 0 | 30 | 0 | 3 | 18 | 100 | 61 | 100 | 0 | 3 |
| G | 9 | 100 | 0 | 0 | 0 | 0 | 0 | 70 | 6 | 0 | 6 | 0 | 0 | 0 | 94 | 0 |
| C | 0 | 0 | 3 | 0 | 0 | 3 | 0 | 0 | 91 | 0 | 6 | 0 | 0 | 0 | 6 | 0 |
| U | 0 | 0 | 15 | 0 | 0 | 91 | 100 | 0 | 3 | 97 | 70 | 0 | 39 | 0 | 0 | 97 |
| Cons. | A*** | G | W | A | A | U | U | R | C | U | W | A | W | A | G | U |

FMN riboswitch, L2-L6 only (1937 sequences)

| | T-loop | | | | | | | J2/3 | | T-loop receptor | | | | | | |
|---|---|---|---|---|---|---|---|---|---|---|---|---|---|---|---|---|
| | 1 | 2 | 3 | 4 | 5 | 6 | 7 | 1 | 2 | 1 | 2 | 3 | 4 | 5 | 6 | 7 |
| A | 0.1 | 0 | 57.5 | 95.3 | 99.8 | 7.2 | 0.1 | 96.0 | 0 | 0 | 6.6 | 100 | 45.0 | 100 | 0 | 0 |
| G | 0.1 | 99.8 | 7.8 | 4.6 | 0.1 | 21.9 | 0 | 4.0 | 0 | 0 | 18.9 | 0 | 7.0 | 0 | 100 | 0.2 |
| C | 15.1 | 0 | 16.4 | 0.1 | 0 | 8.5 | 0.9 | 0 | 96.5 | 0.1 | 25.1 | 0 | 21.4 | 0 | 0 | 0.1 |
| U | 84.7 | 0.2 | 18.4 | 0.1 | 0.2 | 62.3 | 99.0 | 0 | 3.5 | 99.9 | 49.4 | 0 | 26.5 | 0 | 0 | 99.8 |
| Cons. | Y | G | H | A | A | N | U | A | C | U | N | A | H | A | G | U |

FMN riboswitch, combined L2-L6 and L3-L5 (2749 sequences total)

| | T-loop | | | | | | | J2/3 | | T-loop receptor | | | | | | |
|---|---|---|---|---|---|---|---|---|---|---|---|---|---|---|---|---|
| | 1 | 2 | 3 | 4 | 5 | 6 | 7 | 1 | 2 | 1 | 2 | 3 | 4 | 5 | 6 | 7 |
| A | >0.1 | 0 | 57.8 | 93.2 | 99.8 | 6.2 | 0.1 | 91.1 | 0 | 0.1 | 8.7 | >99.9 | 37.7 | 100 | 0.1 | 0 |
| G | 0.1 | 99.5 | 12.7 | 6.4 | 0.1 | 15.7 | 0 | 8.9 | 0 | 0 | 24.5 | 0 | 5.4 | 0 | 99.9 | 0.2 |
| C | 10.9 | 0.3 | 13.1 | <0.1 | 0 | 10.7 | 1.4 | 0 | 97.3 | <0.1 | 18.5 | 0 | 27.7 | 0 | 0 | 5.6 |
| U | 88.9 | 0.2 | 16.4 | 0.3 | 0.1 | 67.3 | 98.5 | 0 | 2.7 | 99.9 | 48.3 | <0.1 | 29.2 | 0 | 0 | 94.2 |
| Cons. | Y*** | G | H | A | A | N | U | A | C | U | N | A | H | A | G | U |

‡Gray boxed nucleotides represent the auxiliary helix.

*Nucleotide position within the element. Positions 1-5 in the T-loop represent the classic T-loop motif, while positions 1-5 in the TLR are the principal receptor motif. Grey shading represents the auxiliary helix.

**These numbers represent the percentage (%) of each nucleotide occupying a position within the alignment.

***Assignment of consensus is based upon the nucleotide occupying the majority at that position (>90% for identity), or shared nucleotide consensus if one nucleotide does not constitute >90% frequency at a given position.

### Table S2. TL/4nt-TLR nucleotide frequencies and consensus

Cobalamin, class II riboswitch (86 sequences)

| | T-loop | | | | | | | J2/3‡ | | T-loop receptor | | | | | | |
|---|---|---|---|---|---|---|---|---|---|---|---|---|---|---|---|---|
| | 1* | 2 | 3 | 4 | 5 | 6 | 7 | 1 | 2 | 1 | --- | 3 | 4 | 5 | 6 | 7 |
| A | 2** | 1 | 83 | 98 | 98 | 7 | 3 | 1 | 0 | 0 | | 99 | 87 | 100 | 0 | 0 |
| G | 0 | 98 | 3 | 2 | 0 | 1 | 0 | 99 | 0 | 0 | | 1 | 3 | 0 | 100 | 0 |
| C | 9 | 0 | 2 | 0 | 0 | 1 | 0 | 0 | 0 | 0 | | 0 | 0 | 0 | 0 | 1 |
| U | 88 | 1 | 12 | 0 | 2 | 91 | 97 | 0 | 100 | 100 | | 0 | 9 | 0 | 0 | 99 |
| Cons. | U | G | W | A | A | U | U | A | C | U | | A | H | A | G | U |

FMN riboswitch, L2-L6 only (1470 sequences)

| | T-loop | | | | | | | J2/3 | | T-loop receptor | | | | | | |
|---|---|---|---|---|---|---|---|---|---|---|---|---|---|---|---|---|
| | 1 | 2 | 3 | 4 | 5 | 6 | 7 | 1 | 2 | 1 | --- | 3 | 4 | 5 | 6 | 7 |
| A | 0.4 | 0 | 53.4 | 84.5 | 98.6 | 5.5 | 0 | 99.7 | 0 | 0 | | 99.9 | 27.9 | 100 | 0.5 | 0 |
| G | 0.3 | 97.3 | 1.4 | 15.4 | 0.3 | 6.7 | 0 | 0.1 | 0 | 0 | | 0.1 | 2.1 | 0 | 99.5 | 0.1 |
| C | 4.6 | 0.1 | 21.4 | 0 | 0 | 1.2 | 0.5 | 0.2 | 92.6 | 0.1 | | 0 | 13.0 | 0 | 0 | 0 |
| U | 94.7 | 2.6 | 23.7 | 0.1 | 1.1 | 86.5 | 99.5 | 0 | 7.4 | 99.9 | | 0 | 57.0 | 0 | 0 | 99.9 |
| Cons. | U*** | G | H | R | A | D | U | A | C | U | | A | H | A | G | U |

FMN riboswitch, combined L2-L6 and L3-L5 (2194 total sequences)

| | T-loop | | | | | | | J2/3 | | T-loop receptor | | | | | | |
|---|---|---|---|---|---|---|---|---|---|---|---|---|---|---|---|---|
| | 1 | 2 | 3 | 4 | 5 | 6 | 7 | 1 | 2 | 1 | --- | 3 | 4 | 5 | 6 | 7 |
| A | 0.4 | 0.1 | 56.6 | 79.6 | 98.9 | 3.8 | 0 | 81.9 | 0 | 0.1 | | 99.6 | 34.1 | 100 | 0.4 | 0 |
| G | 0.2 | 94.0 | 1.5 | 20.1 | 0.2 | 4.8 | 0 | 18.0 | 0.9 | 0 | | 0.1 | 2.6 | 0 | 98.7 | 0.2 |
| C | 3.4 | 0.2 | 18.2 | 0.2 | 0 | 7.1 | 0.8 | 0.1 | 94.1 | 0.1 | | 0.2 | 12.4 | 0 | 0.9 | 17.8 |
| U | 96.0 | 5.7 | 23.7 | 0.1 | 0.9 | 84.3 | 99.2 | 0 | 5.0 | 99.9 | | 0.1 | 50.9 | 0 | 0 | 82.0 |
| Cons. | U | G | H | R | A | Y | U | A | C | U | | A | H | A | G | U |

‡Gray boxed nucleotides represent the auxiliary helix.

*Nucleotide position within the element. Positions 1-5 in the T-loop represent the classic T-loop motif, while positions 1-5 in the TLR are the principal receptor motif. Grey shading represents the auxiliary helix.

**These numbers represent the percentage (%) of each nucleotide occupying a position within the alignment.

***Assignment of consensus is based upon the nucleotide occupying the majority at that position (>90% for identity), or shared nucleotide consensus if one nucleotide does not constitute >90% frequency at a given position.

### Table S3. TL/IL-TLR nucleotide frequencies and consensus

Cobalamin, class II, 19 sequences

| | T-loop | | | | | | | J4/5‡ | | 5' IL receptor | | | 3' IL receptor | | | |
|---|---|---|---|---|---|---|---|---|---|---|---|---|---|---|---|---|
| | 1* | 2 | 3 | 4 | 5 | 6 | 7 | 1 | 2 | 1 | 2 | 3 | 1 | 2 | 3 | 4 |
| A | 0** | 0 | 53 | 79 | 100 | 53 | 0 | 74 | 0 | 0 | 79 | 79 | 74 | 100 | 0 | 0 |
| G | 0 | 100 | 5 | 21 | 0 | 0 | 0 | 26 | 0 | 0 | 21 | 0 | 16 | 0 | 100 | 0 |
| C | 0 | 0 | 16 | 0 | 0 | 0 | 0 | 0 | 100 | 0 | 0 | 0 | 0 | 0 | 0 | 0 |
| U | 100 | 0 | 26 | 0 | 0 | 47 | 100 | 0 | 0 | 100 | 0 | 21 | 11 | 0 | 0 | 100 |
| Cons. | U*** | G | H | R | A | W | U | R | C | U | R | W | D | A | G | U |

Cobalamin, class 1, 7469 sequences

| | T-loop | | | | | | | J4/5 | | 5' IL receptor | | | 3' IL receptor | | | |
|---|---|---|---|---|---|---|---|---|---|---|---|---|---|---|---|---|
| | 1 | 2 | 3 | 4 | 5 | 6 | 7 | 1 | 2 | 1 | 2 | 3 | 1 | 2 | 3 | 4 |
| A | 0.2 | 0.9 | 43.1 | 59.5 | 99.2 | 63.6 | 9.5 | 50.4 | 0 | 0.1 | 64.8 | 85.2 | 49.1 | 99.4 | 0.3 | 0.1 |
| G | 0.4 | 86.3 | 8.0 | 39.3 | 0.6 | 16.9 | 9.1 | 49.5 | 0.1 | <0.1 | 30.9 | 2.0 | 43.8 | 0.2 | 99.5 | 0.2 |
| C | 0.9 | 1.8 | 27.7 | 0.7 | 0.2 | 4.0 | 2.8 | 0.1 | 99.0 | 0.2 | 3.4 | 0.9 | 5.1 | 0.3 | 0.1 | 44.2 |
| U | 98.6 | 11.0 | 21.3 | 0.6 | 0.1 | 15.5 | 78.6 | 0.1 | 0.9 | 99.7 | 0.9 | 11.9 | 2.0 | 0.1 | 0.1 | 55.5 |
| Cons. | U | K | H | R | A | D | U | R | C | U | R | W | R | A | G | Y |

FMN, L2 – L5, 1807 sequences

| | T-loop (L5) | | | | | | | J5/6 | | 5' IL receptor | | | 3' IL receptor | | | |
|---|---|---|---|---|---|---|---|---|---|---|---|---|---|---|---|---|
| | 1 | 2 | 3 | 4 | 5 | 6 | 7 | 1 | 2 | 1 | 2 | 3 | 1 | 2 | 3 | 4 |
| A | 0 | 0.5 | 43.6 | 40.0 | 99.9 | 40.0 | 2.5 | 18.9 | 0 | 0.2 | 64.0 | 73.4 | 47.7 | 99.9 | 0.1 | 0.1 |
| G | 0 | 89.1 | 3.8 | 58.9 | 0.1 | 58.9 | 21.0 | 81.1 | 0.1 | 0 | 35.6 | 6.0 | 27.7 | 0.1 | 99.8 | 0.1 |
| C | 0.2 | 5.2 | 26.6 | 0.94 | 0 | 0.9 | 4.8 | 0 | 99.9 | 0 | 0.3 | 0 | 12.6 | 0.1 | 0.1 | 75.9 |
| U | 99.8 | 5.2 | 26.1 | 0.11 | 0 | 0.1 | 71.7 | 0 | 0 | 99.8 | 0.1 | 20.6 | 12.0 | 0 | 0 | 24.0 |
| Cons. | U | B | H | R | A | R | G/U | R | C | U | R | W | R | A | G | Y |

‡Gray boxed nucleotides represent the auxiliary helix.

*Nucleotide position within the element. Positions 1-5 in the T-loop represent the classic T-loop motif, while positions 1-5 in the TLR are the principal receptor motif. Grey shading represents the auxiliary helix.

**These numbers represent the percentage (%) of each nucleotide occupying a position within the alignment.

***Assignment of consensus is based upon the nucleotide occupying the majority at that position (>90% for identity), or shared nucleotide consensus if one nucleotide does not constitute >90% frequency at a given position.

**Table S4. Data collection statistics for each screen.**

| Library | nt varied | observed[1] | unique | functional[2] | prob.[3] |
|---|---|---|---|---|---|
| TL | 7 | 53300 | 57 | 45 | 96% |
| 5nt-TLR | 7 | 49700 | 75 | 59 | 95 |
| 4nt-TLR | 6 | 18200 | 36 | 21 | 99 |
| IL | 7 | 67100 | 66 | 55 | 98 |
| Aux_helix | 4 | 760 | 21 | 16 | 95 |

[1]Number of total colonies screened, rounded to the nearest 100.
[2]Colonies that were showed better that 2-fold Cbl-dependent repression
[3]Probability of observing any individual sequence within the library as calculated using Equation 1 from Reetz, Kahakeaw and Lohmer.[1]

**Table S5. Biological variation in the auxiliary helix.**

| Pairing | FMN, 5nt-TLR | FMN, 4nt-TLR | FMN, IL-TLR | Cbl IL-TLR |
|---|---|---|---|---|
| **AC/GU** | 2431 | 1359 | 338 | 3607 |
| **GC/GC** | 147 | 0 | 1371 | 3153 |
| **GC/GU** | 90 | 1 | 93 | 488 |
| **AU/GU** | 68 | 100 | 0 | 28 |
| **GU/AC** | 4 | 0 | 0 | 0 |
| **AC/GG** | 3 | 0 | 0 | 0 |
| **GC/GG** | 3 | 0 | 0 | 0 |
| **AC/GC** | 2 | 0 | 0 | 0 |
| **GU/GC** | 1 | 0 | 0 | 31 |
| **AU/AU** | 0 | 7 | 0 | 0 |
| **AG/CU*** ** | 0 | 0 | 2 | 2 |
| **Total** | 2749 | 1467 | 1804 | 7309 |

**Table S6. DNA oligonucleotides used in this study**

| DNA Oligonucleotide | Sequence (5′ to 3′) |
|---|---|
| pRR_forward | GCGCTAGCCACAGCTAACAC |
| 5′_library1 (TL-screen) | CACGACATGCATAAGGCTCGTATAATATATTCATATAATAATGGCCTAAAAGCGTAGTGGGAAAGTGACGNNNNNTTCGTCCAGATTACTTGATACGG |
| 3′_library1 | ACAGGAAAGCTTGGCGTAATCATCAGCATGTTGAGTCTCCTTGCTCTGTATGTTGTATGTTGTATGGCCTAGGTGGCATTCGGAGTANNACCGTATCAAGTAATCTGGACG |
| 5′_library2 (5nt-TLR screen) | CACGACATGCATAAGGCTCGTATAATATATTCATATAATAATGGCCTAAAAGCGTAGTGGGAAAGTGACGTGAAANNCGTCCAGATTACTTGATACGG |
| 3′_library2 | ACAGGAAAGCTTGGCGTAATCATCAGCATGTTGAGTCTCCTTGCTCTGTATGTTGTATGTTGTATGGCCTAGGTGGCATTCGGAGNNNNNCCGTATCAAGTAATCTGGACG |
| 5′_library3 (4nt-TLR screen) | Same as 5′_library2 |
| 3′_library3 | ACAGGAAAGCTTGGCGTAATCATCAGCATGTTGAGTCTCCTTGCTCTGTATGTTGTATGTTGTATGGCCTAGGTGGCATTCGGAGNNNNCCGTATCAAGTAATCTGGACG |
| 5′_library4 (IL-TLR screen) | CACGACATGCATAAGGCTCGTATAATATATTCATATAATAATGGCCTAAAAGCGTAGTGGGAAACAATGTGAAANNCATTGACATTACTTGATACGGNNNGCGCTTCGGCGCNNGTCCGAATGCCACCTAGGCC |
| 3′_library4 | ACAGGAAAGCTTGGCGTAATCATCAGCATGTTGAGTCTCCTTGCTCTGTATGTTGTATGTTGTATGGCCTAGGTGGCATTCGGAC |
| 5′_library5 (BP screen) | CACGACATGCATAAGGCTCGTATAATATATTCATATAATAATGGCCTAAAAGCGTAGTGGGAAAGTGACGTGAAATTCGTCCNNATTACTTGATACGGTTATA |
| 3′_library5 | ACAGGAAAGCTTGGCGTAATCATCAGCATGTTGAGTCTCCTTGCTCTGTATGTTGTATGTTGTATGGCCTAGGTGGCATTCGGNNTATAACCGTATCAAGTAAT |
| 5′_env8_forward | TTTACGGGCATGCATAAGGCTCGTATA |
| Env8_A | ATGCATAAGGCTCGTATAATATATTCATATAATAATGGCCTAAAAGCGTAGTGGGAAAGT |
| Env8_B1 | CCGTATCAAGTAATCTGGACGAATTTCACGTCACTTTCCCACTACGCTTTTAGG |
| Env8_B2 | CCGTATCAAGTAATCTGGACGAATTTCGCGTCACTTTCCCACTACGCTTTTAGG |
| Env8_B3 | CCGTATCAAGTAATCTGGACGAATTACACGTCACTTTCCCACTACGCTTTTAGG |
| Env8_B4 | CCGTATCAAGTAATCTGGACGAATTGCACGTCACTTTCCCACTACGCTTTTAGG |
| Env8_B5 | CCGTATCAAGTAATCTGGACGAATTCCACGTCACTTTCCCACTACGCTTTTAGG |

| | |
|---|---|
| Env8_B6 | CCGTATCAAGTAATCTGGACGAAGCACCCGTCACTTTCCCACTACGCTTTTAGG |
| Env8_C1 | CGTCCAGATTACTTGATACGGTTATACTCCGAATGCCACCTAGGCCATACAACATACAAC |
| Env8_C2 | CGTCCAGATTACTTGATACGGTAATACTCCGAATGCCACCTAGGCCATACAACATACAAC |
| Env8_C3 | CGTCCAGATTACTTGATACGGTCATACTCCGAATGCCACCTAGGCCATACAACATACAAC |
| Env8_C4 | CGTCCAGATTACTTGATACGGTGATACTCCGAATGCCACCTAGGCCATACAACATACAAC |
| Env8_C5 | CGTCCAGATTACTTGATACGGTGGTACTCCGAATGCCACCTAGGCCATACAACATACAAC |
| Env8_D | CTTGGCGTAATCATCAGCATGTTGAGTCTCCTTGCTCTGTATGTTGTATGTTGTATGGCC |
| 3'_Env8 | GGCATGCAAGCTTGGCGTAATCATCAGCATG |
| Env8_T10_B1 | CCGTATCAAGTAATGTGGACGAATTTCACGTCACTTTCCCACTACGCTTTTAGG |
| Env8_T10_B2 | CCGTATCAAGTAATGTGGACGACTTTCACGTCACTTTCCCACTACGCTTTTAGG |
| Env8_T10_B3 | CCGTATCAAGTAATGTGGACGAATTACACGTCACTTTCCCACTACGCTTTTAGG |
| Env8_T10_B4 | CCGTATCAAGTAATGTGGACGAATTTCGCGTCACTTTCCCACTACGCTTTTAGG |
| Env8_T10_B5 | CCGTATCAAGTAATGTGGACGAATTGCACGTCACTTTCCCACTACGCTTTTAGG |
| Env8_T10_C1 | CGTCCACATTACTTGATACGGTTATAGTCCGAATGCCACCTAGGCCATACAACATACAAC |
| Env8_T10_C2 | CGTCCACATTACTTGATACGGTTACAGTCCGAATGCCACCTAGGCCATACAACATACAAC |
| Env8_T10_C3 | CGTCCACATTACTTGATACGGTTAAAGTCCGAATGCCACCTAGGCCATACAACATACAAC |
| Env8_T10_C4 | CGTCCACATTACTTGATACGGTGAAAGTCCGAATGCCACCTAGGCCATACAACATACAAC |
| Env8_T10_C5 | CGTCCACATTACTTGATACGGTGATAGTCCGAATGCCACCTAGGCCATACAACATACAAC |
| Env8_T10_C6 | CGTCCACATTACTTGATACGGTCATAGTCCGAATGCCACCTAGGCCATACAACATACAAC |

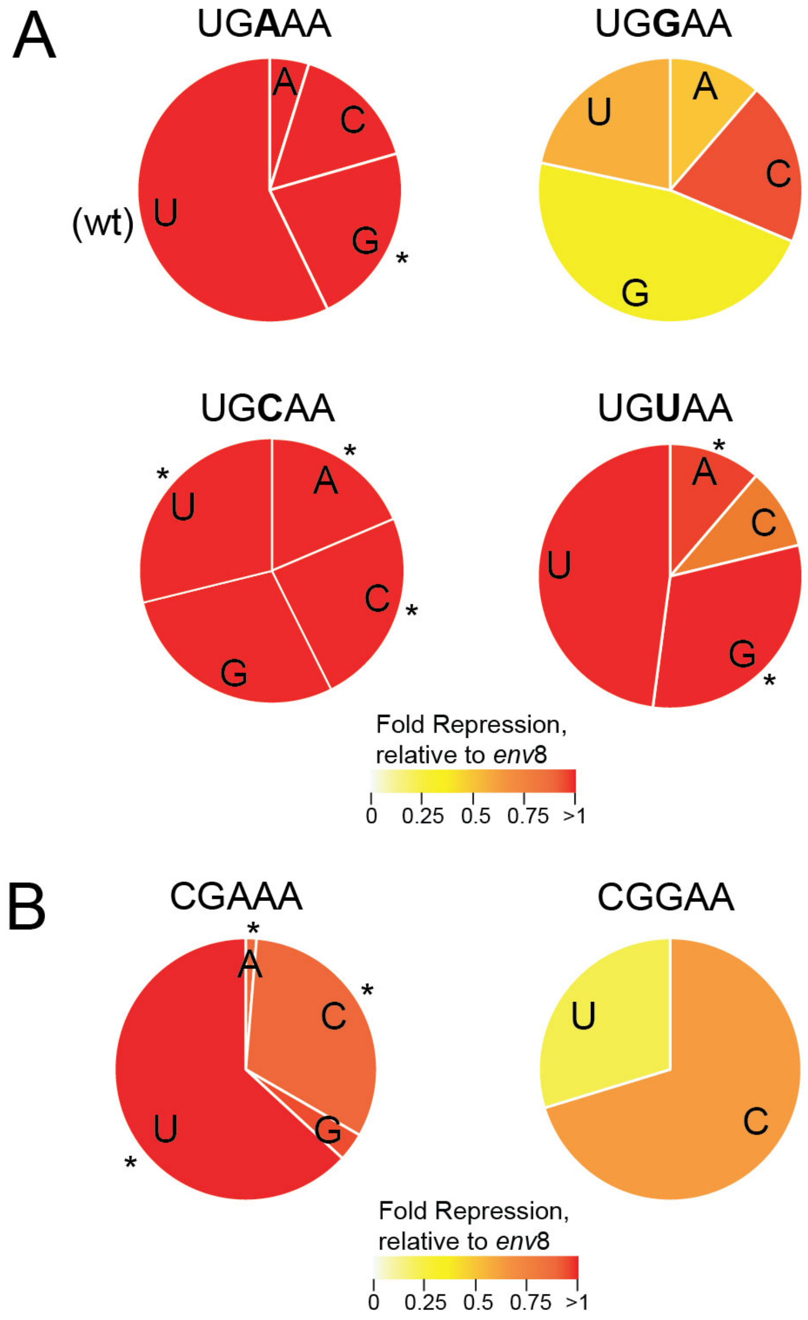


**Figure S1. Nucleotide preferences for nucleotides of the 5nt-TLR interacting with the T-loop.** (A) Population and activity of the 5nt-TLR nucleotide interaction with position 3 of the UGNAA T-loops (contact *i,* **Figure 1(e)**). (B) Population and activity of the 5nt-TLR interacting with the CGRAA T-loops (contact *i,* **Figure 1(e)**).

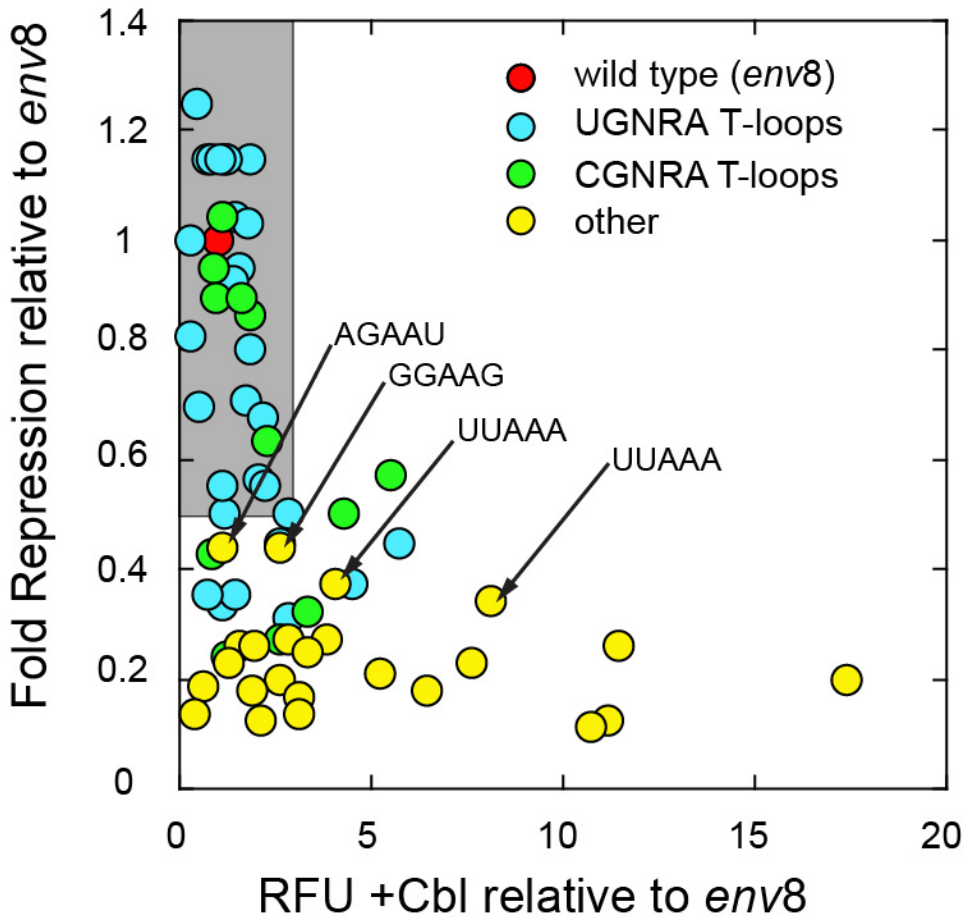


**Figure S2. Distribution of most prevalent sequence motifs in the T-loop genetic screen.** The plot is the same as shown in Figure 2D but data points are colored to reflect the sequence motif. Note that the majority of the variants that fall into the UNANA and CNANA sequence motifs (cyan and green, respectively) cluster in the region of the plot representing moderate-to-highly functional sequences. Rare sequences or those not represented in the FMN riboswitches (yellow) are less functional or non-functional. Highlighted is a T-loop found in the tRNA family (UUAAA; note that the two data points represent different sequences in the TLR) and ribosomal RNA (GGAAG), indicating that these sequences, which are outside the FMN/Cbl T-loop consensus, are still functional. Sequence variant AGAAU represents a sequence in which the U-A base pair between positions 1 and 5 is reversed and likely forms a genetic Watson Crick pair.

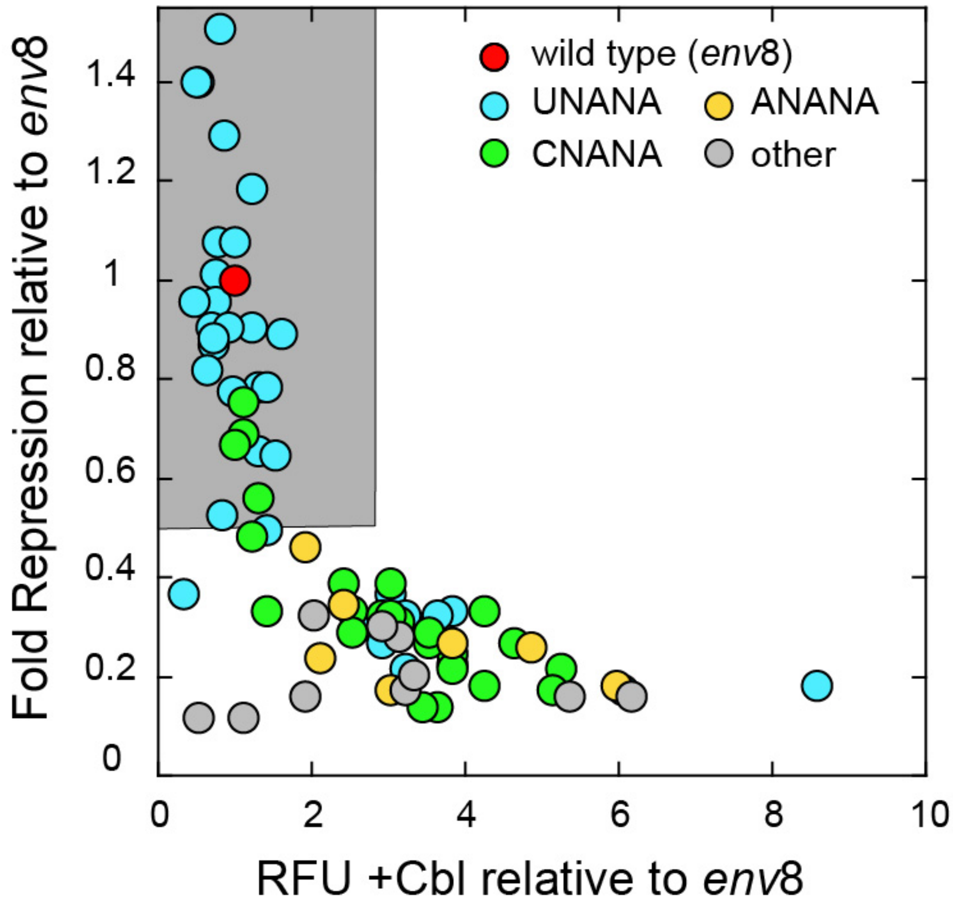


**Figure S3. Distribution of most prevalent sequence motifs in the 5nt-TLR genetic screen.** The plot is the same as shown in Figure 3D but data points are colored to reflect the sequence motif. Note that all of the variants that fall into the UNANA and CNANA sequence motifs (cyan and green, respectively) cluster in the region of the plot representing moderate-to-highly functional sequences.

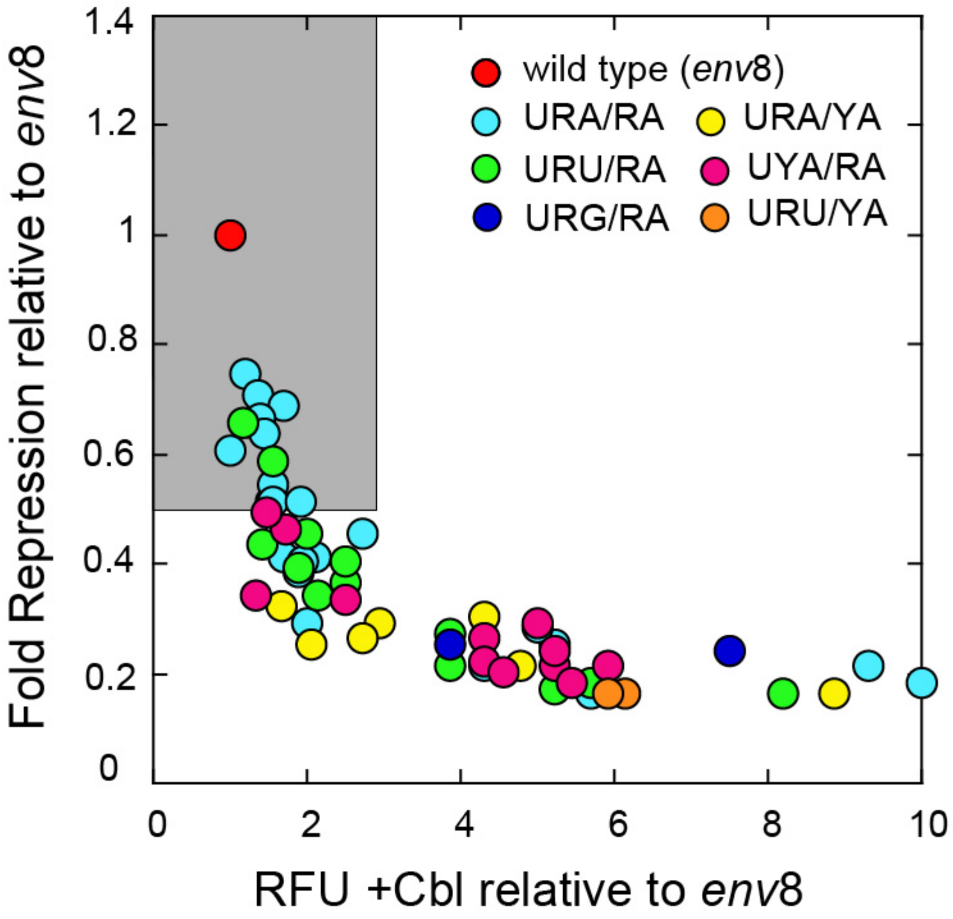


**Figure S4. Distribution of most prevalent sequence motifs in the IL-TLR genetic screen.** The plot is the same as shown in Figure 5F but data points colored to reflect the different sequence motifs.
