## Supplementary material for "Context-dependence of T-loop mediated long-range RNA tertiary interactions": FASTA_files_README.docx

Contents of Folder:

**Phylogenetic Alignments:**

1. B12_C1_ILTLR.fa: Fasta format file containing alignment of all T-loop/TLR interactions in class-I cobalamin riboswitches, taken from Rfam accession number RF00174 (Rfam 14.0). Sequences have been truncated to include L4 (T-loop), J4/5 (5’-side of auxiliary helix), J6/7 (5’-side of IL-TLR) and J7/6 (3’-side of IL-TLR) in the format XXXXXXX-XX-XXX-XXXX corresponding to each region, respectively. Sequence accession numbers have been preserved to enable acquisition of the entire RNA sequence from the Rfam database.

2. B12_C2_4ntTLR.fa: Fasta format file containing alignment of all T-loop/4nt-TLR interactions from class-II cobalamin riboswitches, taken from Rfam accession number RF01689 (Rfam 14.0). Sequences have been truncated to include L4 (T-loop), J4/5 (5’-side of auxiliary helix), and L6 (4ntTLR) in the format XXXXXXX-XX-XXXXXX corresponding to each region, respectively. Sequence accession numbers have been preserved to enable acquisition of the entire RNA sequence from the Rfam database.

3. B12_C2_5ntTLR.fa: Fasta format file containing alignment of all T-loop/5nt-TLR interactions from class-II cobalamin riboswitches, taken from Rfam accession number RF01689 (Rfam 14.0). Sequences have been truncated to include L4 (T-loop), J4/5 (5’-side of auxiliary helix), and L6 (4ntTLR) in the format XXXXXXX-XX-XXXXXXX corresponding to each region, respectively. Sequence accession numbers have been preserved to enable acquisition of the entire RNA sequence from the Rfam database.

4. B12_C2_ILTLR.fa: Fasta format file containing alignment of all T-loop/IL-TLR interactions from class-II cobalamin riboswitches, taken from Rfam accession number RF01689 (Rfam 14.0). Sequences have been truncated to include L4 (T-loop), J4/5 (5’-side of auxiliary helix), J6/7 (5’-side of IL-TLR) and J7/6 (3’-side of IL-TLR) in the format XXXXXXX-XX-XXX-XXXX corresponding to each region, respectively. Sequence accession numbers have been preserved to enable acquisition of the entire RNA sequence from the Rfam database.

5. FMN_4ntTLR_merged.fa: Fasta format file containing alignment of all T-loop/4nt-TLR interactions from class-II cobalamin riboswitches, taken from Rfam accession number RF01689 (Rfam 14.0). Sequences have been truncated to include L2 or L5 (T-loop), J2/3 or J5/6 (5’-side of auxiliary helix), and L6 or L3 (4ntTLR) in the format XXXXXXX-XX-XXXXXX corresponding to each region, respectively. Note that in this file the L2-L6 and L3-L5 interactions have been merged after permuting the L3-L5 sequences to align with the L2-L6 sequences. Sequence accession numbers have been preserved to enable acquisition of the entire RNA sequence from the Rfam database.

6. FMN_5ntTLR_merged.fa: Fasta format file containing alignment of all T-loop/4nt-TLR interactions from class-II cobalamin riboswitches, taken from Rfam accession number RF01689 (Rfam 14.0). Sequences have been truncated to include L2 or L5 (T-loop), J2/3 or J5/6 (5’-side of auxiliary helix), and L6 or L3 (5ntTLR) in the format XXXXXXX-XX-XXXXXXX corresponding to each region, respectively. Note that in this file the L2-L6 and L3-L5 interactions have been merged after permuting the L3-L5 sequences to align with the L2-L6 sequences. Sequence accession numbers have been preserved to enable acquisition of the entire RNA sequence from the Rfam database.

7. FMN_ILTLR.fa: Fasta format file containing alignment of all T-loop/TLR interactions in FMN riboswitches, taken from Rfam accession number RF00050 (Rfam 14.0). Sequences have been truncated to include L5 (T-loop), J5/6 (5’-side of auxiliary helix), J3a/3b (5’-side of IL-TLR) and J3b/3a (3’-side of IL-TLR) in the format XXXXXXX-XX-XXX-XXXX corresponding to each region, respectively. Sequence accession numbers have been preserved to enable acquisition of the entire RNA sequence from the Rfam database.

**Genetic Screen Alignments:**

1. TL_FR.fa: sequences from the T-loop library with fold repression normalized to env8 reported, only those with a FR greater than or equal to 2.0 were included

2. 5ntTLR_FR.fa: sequences from the 5nt T-loop receptor library with fold repression normalized to env8 reported, only those with a FR greater than or equal to 2.0 were included

3. 4ntTLR_FR.fa: sequences from the 4nt T-loop receptor library with fold repression normalized to env8 reported, only those with a FR greater than or equal to 2.0 were included

4. IL_FR.fa: sequences from the Internal Loop T-loop receptor library with fold repression normalized to env8 reported, only those with a FR greater than or equal to 2.0 were included

5. aux_helix_FR.fa: sequences from the auxiliary helix library with fold repression normalized to env8 reported, only those with a FR greater than or equal to 2.0 were included

FASTA files were created by converting the sheets from “Tloop_data_table.xlsx” into .csv and using:

awk -F , '{print ">"$1"_"$4"\n"$9}' {file}.csv > {file}.fas

The information indexed can vary. The above example creates a fasta file with the fluorescence in the presence of cobalamin ($4) but that information can be varied or removed.

**Activity Assay data**

1. Tloop_assay_values.xlsx: Values from activity assays. Each sheet includes all assay data for a given library. Within each sheet, columns denote unique sequences with names assigned based on coordinates of gridded plate in each round. Each value is a ratio of the fluorescence measurement and the optical density at 600 nm. Column 1 denotes whether the value is in absence of ligand (-), presence of ligand (+), or the fold induction (FI) ration. Each unique sequence has at least 9 values for 9 unique measurements, since many have more, those cells that were left empty contain “-2” to be removed during analysis. Analysis of these data completed by converting each sheet into a .csv file to run in Tloop_project_analysis.Rmd.

2. Tloop_data_table.xlsx: Compiled results from analysis via Tloop_project_analysis.Rmd. Each library has two sheets, one for all data values “X_all” and one with only colonies with FI >= 2.0 “X”. Each sheet includes the colony name (rows), median OD corrected fluorescence in presence of CNCbl (col B) absence of CNCbl (col D), the standard error of each (col C and E) and the median fold induction (col F). Additional data was added to output files in col F-L. Those median values are also reported as env8 corrected values (col G-I) and important sequence elements and the full sequence was reported in the neighboring columns.
